# Limited proteolysis-coupled mass spectrometry captures proteome-wide protein structural alterations and biomolecular condensation in living cells

**DOI:** 10.1101/2025.02.21.639445

**Authors:** Franziska Elsässer, Roberta Florea, Felix Räsch, Karsten Weis, Natalie de Souza, Paola Picotti

## Abstract

The function of a protein is determined by its structure, which may change dynamically in response to post-translational modifications, interaction with other molecules, or environmental factors like temperature. Limited proteolysis-coupled mass spectrometry (LiP-MS) captures such structural alterations on a proteome-wide scale via the detection of altered protease susceptibility patterns of proteins. However, this technique has so far required cell lysis, which exposes proteins to non-native conditions and can disrupt labile interactions such as those occurring within biomolecular condensates. To study protein structures directly within cells, we developed in-cell LiP-MS. We optimized conditions for introduction of proteinase K into human cells using electroporation and validated that intracellular cleavage occurs. In-cell LiP-MS captured not only the known binding of rapamycin to FKBP1A within the cell, but also downstream effects of pathway activation. Moreover, it detected global protein structural alterations upon sodium arsenite treatment and captured the structural dynamics of hundreds of proteins from biomolecular condensates with peptide level resolution and within live human cells. We detected known and novel structural alterations of proteins from stress granules as well as from nuclear speckles and validated alteration of nuclear speckles by fluorescence microscopy. Our dataset provides a resource describing the structural changes of human proteins in response to cellular stress and pinpoints structurally altered regions. Further, comparison of LiP-based structural fingerprints before and after cell lysis revealed which human proteins are susceptible to structural change upon cell lysis, therefore guiding the design of future experiments requiring native protein structures.

## Introduction

Proteins serve many functions in cells, regulating growth, differentiation, and survival. Since protein structure is a key determinant of protein function, the study of protein structure before and after various perturbations (e.g., stress, disease, or treatment with drugs) yields insight into the understanding of protein function and dysfunction in health and disease. Structural predictions based on Alphafold or RoseTTAFold are now available for over 98% of human proteins (Baek et al., 2021; Tunyasuvunakool et al., 2021), complementing the large body of experimentally solved protein structures. However, most available protein structures are a static snapshot, and information on protein structural changes between conditions is very limited.

Techniques to determine high-resolution protein structures typically require that the proteins or protein complexes of interest are purified. Such *in vitro* analyses do not recapitulate important properties of the cellular environment such as molecular crowding, presence of small-molecule or macromolecule interactors, and subcellular gradients of pH or metabolites. Also, certain features of protein structure depend on subcellular localization and might be altered upon cell lysis. Low-affinity or dynamic interactions, such as those within biomolecular condensates, can be lost or difficult to capture upon cell lysis also due to dilution of the cellular content (Jain et al., 2016). It is therefore important to study protein dynamics within the native cellular environment.

In-cell nuclear magnetic resonance (NMR) and Förster resonance energy transfer (FRET) techniques provide information on the dynamics of protein structures with atomic resolution inside cells (Ha et al., 1999; Sakakibara et al., 2009), but typically one protein is studied at a time, both techniques require labeling, and FRET requires extensive optimization. Cryo-electron tomography can be used to solve protein structures in their cellular context, but studying protein dynamics remains challenging, and the approach is not applicable on a proteome-wide scale.

We previously developed limited proteolysis coupled mass spectrometry (LiP-MS), a structural proteomics approach that allows the study of structural changes in thousands of proteins simultaneously with peptide-level resolution (Feng et al., 2014). In LiP-MS, native proteins are cleaved by a non-specific protease for a short period of time, followed by trypsin digestion under denaturing conditions, which results in structure-specific peptide patterns that can be quantitatively read out by mass spectrometry (Figure 1a). We have shown that LiP-MS can detect numerous molecular events, including allosteric regulation, changes in enzyme activity, protein-protein interactions, post-translational modification, protein-small molecule interactions, and protein aggregation events(Cappelletti et al., 2021; Piazza et al., 2018). LiP-MS has been applied for drug target identification (Piazza et al., 2020) and to characterize structural changes in biofluids of individuals suffering from Parkinson’s disease (Mackmull et al., 2022). Although powerful, LiP-MS has so far only been applicable to cell or tissue lysates (Malinovska et al., 2023; Schopper et al., 2017), where various protein structures may be altered due to the loss of intracellular conditions.

**Figure 1:**
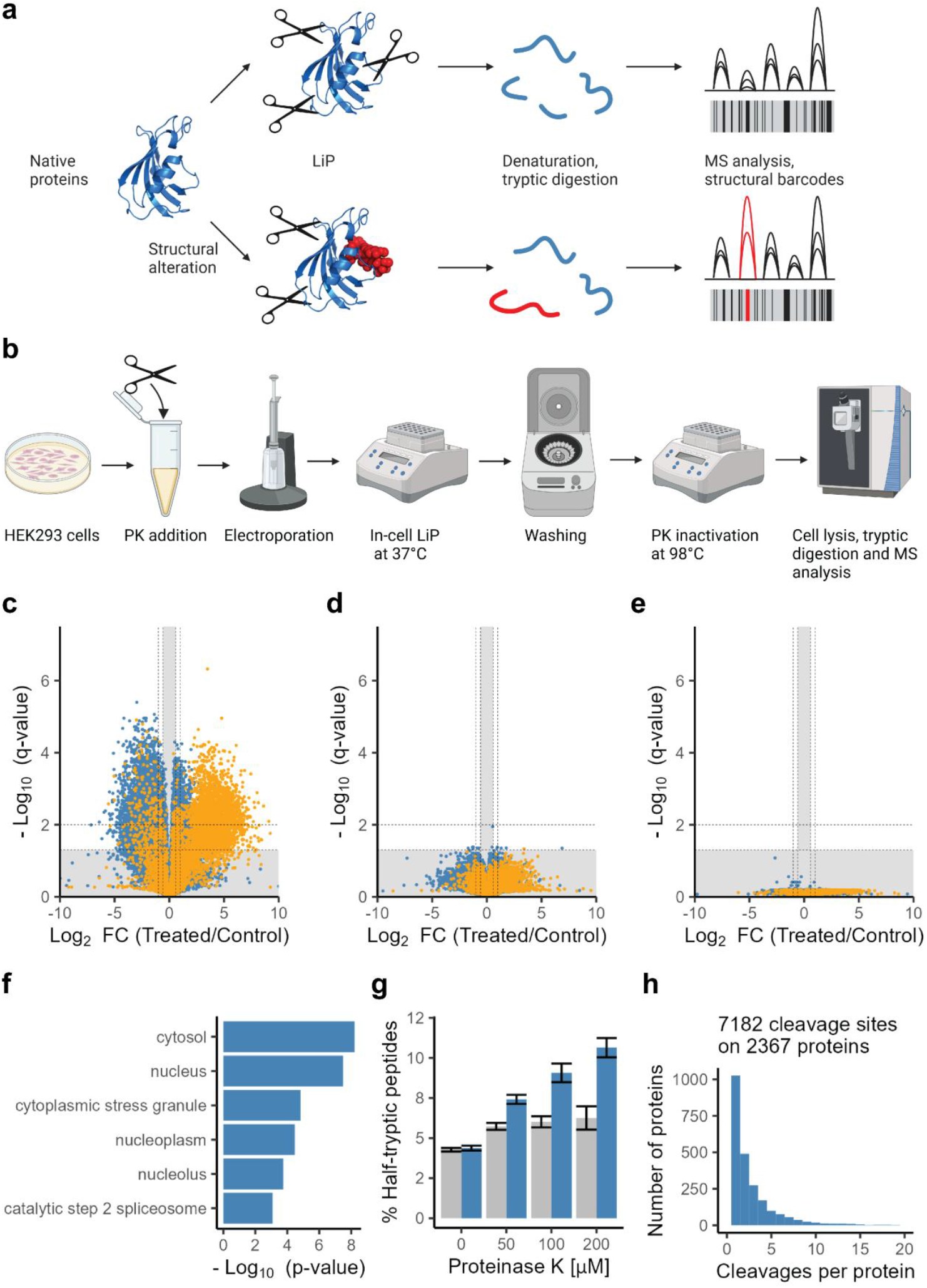
Development of in-cell limited proteolysis-coupled mass spectrometry (in-cell LiP-MS). (**a**) Illustration of LiP-MS. Native proteins are incubated with PK for a short time to generate structure-specific peptides. The samples are then subjected to denaturing conditions, and proteins are digested with trypsin and analyzed by mass spectrometry. A structural change leads to altered proteolytic cleavage by PK. (**b**) Schematic of the in-cell LiP-MS workflow. Low-specificity PK is introduced into cells by electroporation. During a brief incubation, structure-specific peptides are generated. After protease inactivation, cell lysis and trypsin treatment, the peptides are analyzed by mass spectrometry. (**c-e**) Changes in peptide intensities in treated samples compared to control samples. Shown are (c) samples treated with 200 μM PK with and without electroporation, (d) samples with and without 200 μM PK in the absence of electroporation, and (e) samples with and without electroporation (1 pulse of 25 ms at 1000 V) in the absence of PK. Each data point represents a single peptide; half-tryptic peptides are shown in orange, and fully tryptic peptides are shown in blue. The x-axes report the log_2_-transformed ratio of peptide intensities. The y-axes report the negative log_10_-transformed q-value, which is the p-value of the t-test that was adjusted for multiple testing. The shaded gray region marks significance levels (FC ≥ 1.5, q-value < 0.05, n = 5 replicates). (**f**) Gene ontology enrichment analysis of proteins that undergo structural changes upon electroporation of PK (p-value < 0.001). (**g**) Fraction of half-tryptic peptides without (control; grey) and with electroporation (blue) of indicated PK concentration. (**h**) Cleavages per protein in optimized in-cell LiP-MS (100 μM PK, 1 pulse of 25 ms at 1000 V, 2 min incubation time, n = 6 replicates).

Here we describe our development of in-cell LiP-MS. The in-cell analysis is enabled by delivery of proteinase K (PK), a low specificity protease, into living mammalian cells using an electroporation approach optimized to ensure rapid (millisecond) and reproducible intracellular uptake of the protease and the desired level of intracellular cleavage. We detected PK-induced LiP signals across the proteome in an electroporation-dependent manner and validated the delivery of active protease into cells using an intracellular sensor based on FRET. In-cell LiP-MS captured the well-characterized and high-affinity interaction of rapamycin with the protein FKBP1A (Heitman et al., 1991) as was previously observed with standard LiP-MS in cell lysates (Piazza et al., 2020). In-cell LiP also revealed dynamics within biomolecular condensates that are formed in part through low-affinity interactions. Under arsenite stress, in-cell LiP-MS captured structural alterations in known and novel stress granule-related proteins as well as in nuclear speckle proteins, which we could validate with fluorescence microscopy. Our study provides the first available resource of *in vivo* proteome-wide protein structural changes during cellular stress, identifies proteins that undergo structural alterations upon cell lysis, and pinpoints structurally altered regions in proteins, including those involved in biomolecular condensation. Further, the in-cell LiP-MS method will allow the study of protein structural changes under any conditions of interest within living cells with peptide-level resolution and on a proteome-wide scale.

## Results

### Development of in-cell LiP-MS

To enable the study of protein structural states in their native cellular environment, we aimed to identify electroporation conditions that reliably introduce sufficient PK into mammalian cells to yield structurally informative cleavages with good coverage of the proteome. We detached HEK293 cells from the culture dish, washed, and resuspended in PBS (Figure 1b). Immediately after addition of PK, cells were subjected to electroporation and then incubated for 1 min at 37 °C to allow limited proteolysis of intracellular proteins. Extracellular PK was removed by washing, PK activity was quenched by addition of a chaotrope and heating to 98 °C, and samples were snap-frozen. Cells were subsequently lysed, trypsin treated, and quantitative, label-free MS analysis was performed.

To assess whether PK had been delivered into cells, we quantified the fully tryptic peptides (FT; i.e., those with two tryptic ends) and the half-tryptic peptides (HT, i.e., those with only a single tryptic end) under various conditions, thus using the proteolytic cleavage itself as a readout of PK delivery. In contrast to trypsin, which cleaves preferably after lysine and arginine residues, PK has low sequence specificity so that a peptide with one or two non-tryptic termini is likely to be generated by PK cleavage. Therefore, an increase in HT peptide intensity and a decrease in FT peptide intensity in a sample sequentially digested with PK and trypsin, relative to a sample digested with trypsin alone, reports on PK cleavage events. We indeed observed evidence of PK cleavage in cells electroporated in the presence of PK compared to cells incubated with PK but not electroporated (Figure 1c). PK cleavages were detected on 2491 proteins. Without electroporation, addition of PK to cells did not lead to significant increased proteolytic cleavage over control (no PK; Figure 1d). That PK indeed enters the cells upon electroporation was further supported by gene ontology enrichment analysis, which showed that proteins with peptide signatures indicative of PK cleavage were significantly enriched in cytosolic and nucleus-associated terms (Figure 1f). Further, there appears to be no substantial leakage of intracellular proteins upon electroporation since the protein content of the cell supernatant was not changed by electroporation (Figure S1a).

In order to identify conditions that result in rapid and reproducible intracellular delivery of PK with minimal electroporation-induced effects on cellular physiology, we examined proteome-wide changes in peptide and protein levels upon electroporation alone. We observed no significant electroporation-induced changes with one 25-ms pulse at 1000 V or with up to two 25-ms pulses at 800 V (Figure 1e, Figure S1b). Moreover, FT and HT peptides were symmetrically distributed in all tested electroporation settings, suggesting that electroporation alone does not cause protein cleavage (Figure S1b, e).

Under higher voltage and with more pulses, we observed a drop in peptide and protein intensity relative to the non-electroporated control (Figure S1b, c), suggesting that high voltage induces protein aggregation or cleavage. Cell viability as assayed with trypan blue staining did not significantly decrease within 5 min after electroporation with one 25-ms pulse of 1000 V (ANOVA q-value < 0.05; Figure S1d), and we chose these conditions for subsequent experiments.

In order to maximize proteome coverage of structurally-informative PK cleavages, we then optimized the enzyme-to-substrate ratio and determined that extracellular PK concentrations of 100 μM for 3 × 10^6^ cells led to the highest number of reproducible intracellular cleavages (Figure 1g). Although more cleaved peptides were detected at higher PK concentrations with up to 17% of peptides being HT at 1000 μM PK (30% half-tryptic peptides is typical in a standard LiP-MS experiment in lysate), the reproducibility of cleavage decreased compared to 100 μM PK (Figure S2a-c), possibly because of over-digestion. Hence, we chose 100 µM extracellular PK for electroporation in subsequent experiments. We further increased the number of reproducible electroporation-dependent PK cleavages by optimizing incubation time and number of replicates (Figure S3a). Using a 2-minute incubation time and six replicates per condition, 17050 peptides on 3616 of 4760 detected proteins changed significantly relative to control (i.e., PK added, no electroporation; Figure S3a), corresponding to 7182 distinct PK cleavage sites on 2367 proteins (Figure 1h; S3a). The coverage in an experiment comparing two perturbed cell states may be higher, since coverage is affected statistically by the fact that several species of HT peptides are not detected in the non-electroporation control; this is expected to be less of an issue when two electroporated samples are being compared. Prolonging limited proteolysis to 5 minutes did not further increase PK cleavage (Figure S3b), and reducing it yielded lower digestion reproducibility. To conclude, we have identified conditions to deliver PK into HEK293 cells to generate reproducible PK-induced cleavages in about 1600 mainly intracellular proteins without unduly compromising cell viability or the cellular proteome.

### Validation of in-cell LiP-MS

To orthogonally test that PK enters cells upon electroporation and is active intracellularly, we used HEK293 cells expressing a FRET-based sensor of protease activity. The sensor consists of ECFP and YPet proteins separated by a short linker sequence (Gray et al., 2010). Upon excitation of ECFP, yellow YPet fluorescence indicates an intact sensor, whereas cleavage of the linker region between the two fluorescent proteins results in loss of FRET signal and therefore an increase in the ECFP fluorescence relative to that of YPet. We first confirmed that the sensor behaves as expected upon addition of PK to a native lysate of the reporter cells. We observed cleavage within minutes of adding PK, and exposing the protease to our electroporation conditions (one 25-ms pulse of 1000 V) did not affect its activity (Figure 2a). Electroporation of PK into cells also resulted in an increase in the ECFP fluorescence relative to that of YPet, confirming that active PK enters the cells and cleaves the reporter protein (Figure 2b). The larger variation we observe in the in-cell context are likely to be due to cell-to-cell variability, for example in sensor expression and PK internalization, as well as technical variability introduced by the electroporation procedure. Neither addition of PK without electroporation nor electroporation alone affected the sensor fluorescence signal (Figure 2b). These data provide orthogonal evidence that electroporation allows the introduction of active PK into mammalian cells.

**Figure 2:**
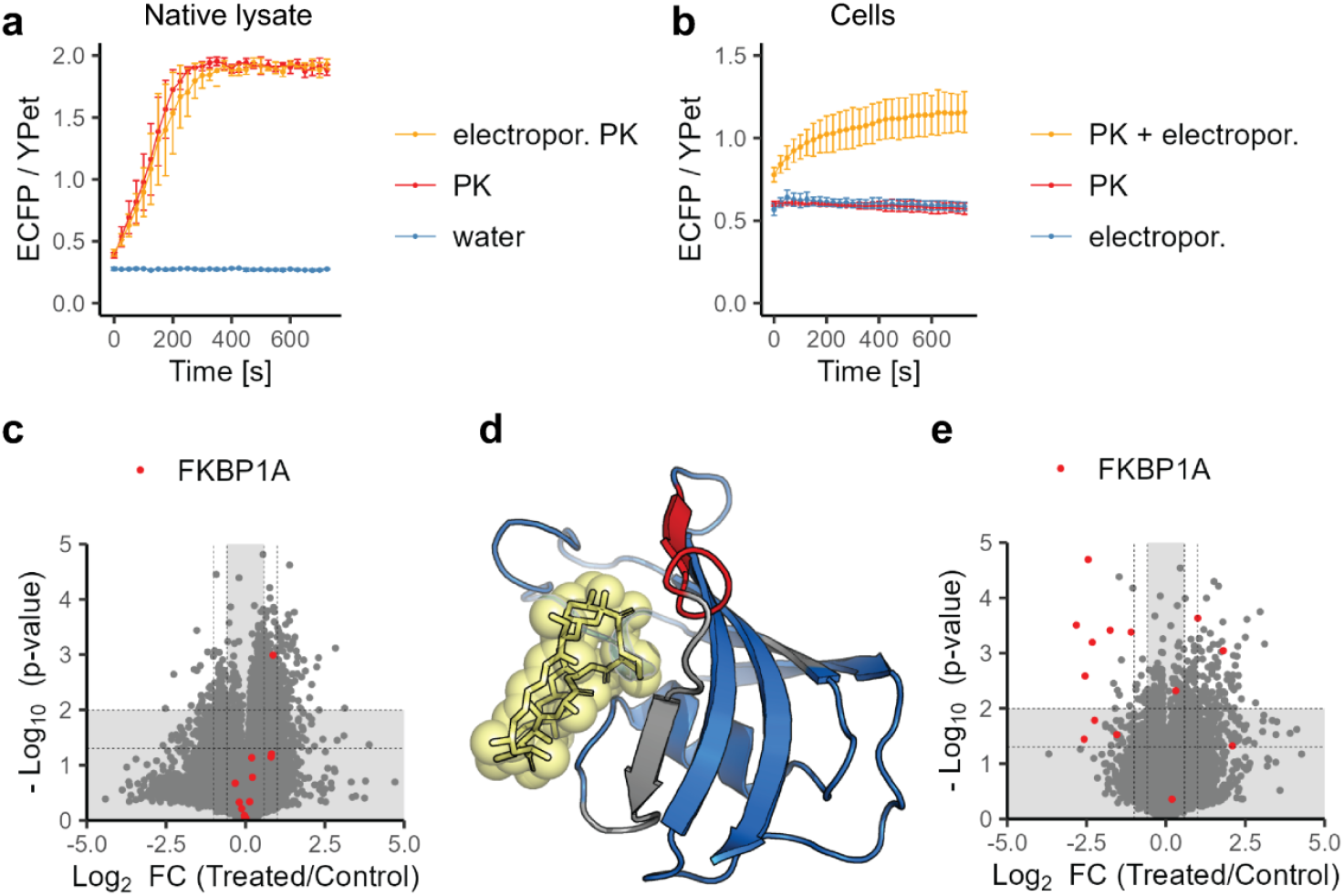
Validation of intracellular PK delivery and target detection by in-cell LiP-MS. (**a**) FRET signal as a function of time of a native lysate of HEK293 cells containing an ECFP/YPET FRET sensor of protease activity that was incubated with PK, PK exposed to electroporation conditions, or water (n = 4 replicates). (**b**) FRET signal from HEK293 cells expressing an ECFP/YPET FRET sensor that were electroporated with PK, electroporated without protease, or that were incubated with PK without electroporation (n = 3 replicates). (**c**) Peptide intensities in HEK293 cells treated with 100 nM rapamycin dissolved in DMSO for 5 min compared to intensities in cells treated only with DMSO in an in-cell LiP setup. Each data point represents a single peptide; peptides of FKBP1A are shown in red. The shaded gray region marks significance levels (FC ≥ 1.5, p-value < 0.01, n = 6 replicates). (**d**) Structure of rapamycin (yellow) and FKBP1A (PDB 2DG3). Regions in blue were detected by mass spectrometry, region in red was significantly changed upon treatment with rapamycin. (**e**) Peptide intensities in a native lysate of HEK293 cells treated with 10 nM rapamycin dissolved in DMSO for 5 min compared to intensities in cells treated only with DMSO (6 replicates). Each data point represents a single peptide; peptides of FKBP1A are shown in red. The shaded gray region marks significance levels (FC ≥ 1.5, p-value < 0.01, n = 4 replicates).

We estimated the number of PK molecules that enter the cell upon electroporation under our chosen experimental conditions by comparison to a calibration curve prepared by analysis of lysates containing four accurately quantified heavy-labeled PK-specific peptides. Cells were treated with PK and subjected to electroporation; control cells were treated with the protease but not electroporated, after which both treated and control cells were thoroughly washed to remove extracellular PK. By comparing to the calibration curve, we calculated that the electroporated sample contained 52.5 pmol PK and that the control sample contained 16.0 pmol PK (Figure S4a). Since the control cells were not electroporated, this is likely to be extracellular PK that was not removed during washing, and should not be able to cleave the intracellular proteome. We reasoned that the difference (36.6 pmol) corresponds to intracellular PK; note that we had initially added 12 nmol of PK to the cells.

Next, we asked whether in-cell LiP-MS can capture known structural alterations associated to high-affinity binding events by making use of the well-characterized binding of rapamycin to the protein FKBP1A. After treatment with 100 nM rapamycin for 5 minutes, cells were electroporated with PK and processed following the previously optimized workflow. We detected an FKPB1A-derived peptide as one of the strongest hits in a comparison of peptide intensities between rapamycin-treated and control (DMSO-treated) cells (Figure 2c; FC > 1.5, p-value < 0.01, n = 6 replicates). Further, this peptide mapped close to the known rapamycin binding site on FKPB1A (Figure 2d). Similar results were obtained for cells treated with 10 μM rapamycin for 10 minutes prior to electroporation of PK (Figure S4c, c; FC > 1.5, q-value < 0.05, n = 6 replicates). Standard LiP-MS of a rapamycin-treated HEK293 cell lysate identified multiple FKBP1A-derived peptides among the most significant hits (FC > 1.5, p-value < 0.01, n = 4 replicates; Figure 2e), as we have previously shown (Piazza et al., 2020).

We observed some variability in the specific peptide that showed altered protease accessibility upon rapamycin treatment. One in-cell LiP detected FKBP1A peptide (Figure S4c, d) also showed changes in the lysate, whereas the other in-cell LiP detected peptide (Figure 2c, d) did not. In both standard and in-cell LiP, several proteins besides FKBP1 showed structural changes after rapamycin treatment (100 nM rapamycin, 5 minutes); we identified 333 (out of 3534 detected) structurally changing proteins with in-cell LiP, and 141 structurally changing proteins (out of 3183 detected) with the standard in-lysate method. Interestingly, the overlap between these sets of proteins was small; only 30 proteins changed structurally in both experiments (Figure S4b) despite good overlap in coverage. This could be because in-lysate LiP may detect target engagement and off-target effects whereas in-cell LiP data may be dominated by effects downstream of the targets. In the in-cell LiP experiment of cells treated with 10 μM rapamycin for 10 minutes, 60 proteins changed structurally (Figure S4b).

The percentage of HT peptides was lower for in-cell LiP than for classical LiP in the native lysate (Figure S4e, f), indicating that there is less PK cleavage in the in-cell context. In this comparison, we used a similar ratio of protein substrate and PK in both conditions: i.e., 100 μg protein and 35 pmol PK (1 μg) in the lysate and 3 million cells (800-1000 μg protein) and 12 nmol (347 μg) PK for in-cell LiP. PK was among the most abundant proteins detected in both the lysate and in cells (Figure S4e, f), suggesting that its amount is not a limiting factor, and the coefficient of variation was similar in both approaches. The lower intracellular cleavage could be the result of lower intracellular diffusion due to crowding and the presence of the cytoskeleton, thus reducing access of PK to the proteome. Moreover, there is likely to be a distribution of PK uptake, with many cells likely internalizing only low amounts or no PK. Taken together, these data show that limited proteolysis in cells captures a known binding event in a complex native background.

Next, we asked whether we could use our data to detect proteins that undergo structural alterations upon cell lysis, reasoning that this set of proteins would benefit in particular from the in-cell setup. To identify these proteins, we compared the DMSO-treated control samples of our parallel in-cell LiP and in-lysate LiP experiments. We found overall a relatively good correlation of peptide intensities (Pearson correlation coefficient ρ = 0.70; Figure 3a), indicating that many proteins retain their structure upon cell lysis at least in these experimental conditions. This was similar for both fully-tryptic (ρ = 0.71) and half-tryptic (ρ = 0.69) peptides (Figure S5a), and for most peptides the ratio of intensities in in-cell LiP to lysate LiP was close to 1 (Figure S5b). Nevertheless, there were several proteins or protein regions with relatively poor correlation (i.e., peptides that are not on the diagonal of the correlation plot) and we investigated these proteins further.

**Figure 3:**
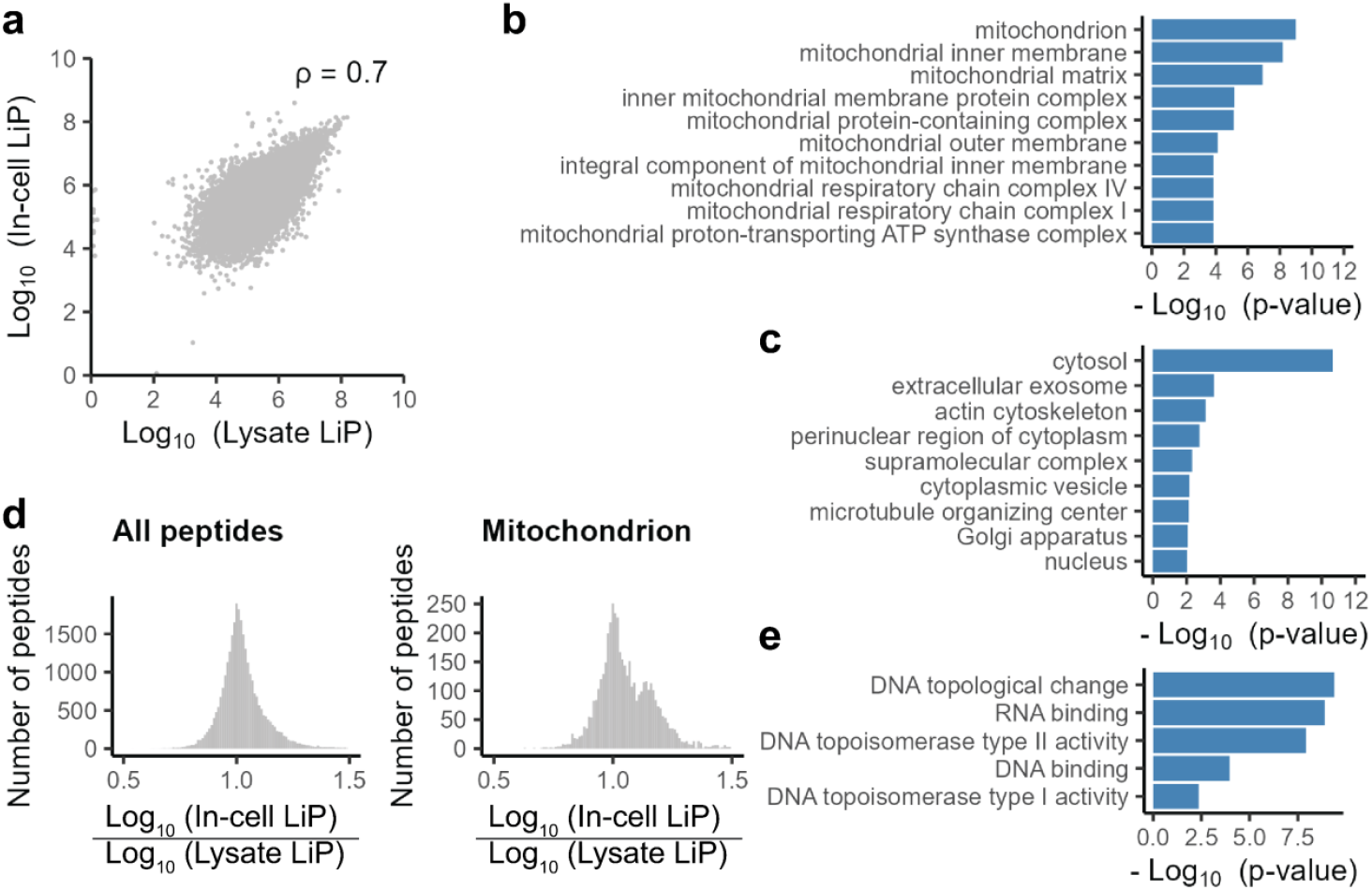
Organelle and domain coverage of LiP-MS in cells and in lysate. (**a**) Correlation of peptide intensities in HEK293 cells between LiP-MS in lysate and in cells after 5 min of DMSO treatment (mean intensity across replicates; in-cell LiP n = 6 replicates; lysate LiP n = 4 replicates); ρ indicates the Pearson correlation coefficient. (**b**,**c**) For proteins with >3 peptides detected in at least 3 replicates of both datasets, a protein score was calculated (protein score: mean ratio of log_10_-transformed peptide intensities between in-cell LiP and lysate LiP across peptides). Gene ontology enrichment of cellular compartments was performed for (b) proteins with protein scores < 0.9 or > 1.1, and (c) protein scores > 0.9 and < 1.1.(p-value < 0.001) (**d**) Ratio of log_10_-transformed peptide intensities between in-cell LiP and lysate LiP, for all peptides and for peptides mapping to proteins located in the mitochondrion based on gene ontology annotation. (**e**) Gene ontology enrichment of protein domains was performed for peptides with a ratio >1.1 of log_10_-transformed peptide intensities between in-cell LiP and lysate LiP (excluding mitochondrial peptides). Domain annotations are from Interpro database (p-value < 0.001).

To test whether peptides with low correlation between in-cell LiP and classical LiP share common features such as a subcellular localization or function, we calculated a protein-level score of similarity between in-cell LiP and classical LiP by averaging the peptide ratios (for proteins >3 peptides). Interestingly, proteins that differ most between the methods (protein score < 0.9 or > 1.1) are located in mitochondria based on the EBI human GOA database (release 2018-02-20; Figure 3b), whereas proteins in the cytosol, nucleus, vesicles and the Golgi apparatus show similar LiP-based structural fingerprints in both methods (protein score > 0.9 and < 1.1; Figure 3c). Correlation at the peptide level was also lower in mitochondria than in other organelles (ρ = 0.65; Figure S5c), and peptide ratios between in-cell LiP and classical LiP revealed two distinct peaks for mitochondrial proteins (Figure 3d). In contrast, peptide ratios of proteins located in other organelles exhibit a single peak with an in-cell:lysate intensity ratio close to 1 (Figure S5d). This indicates two populations of mitochondrial proteins that are differentially cleaved by PK. Since most mitochondrial proteins are encoded in the nucleus, translated in the cytosol, and post-translationally translocated into the mitochondrial compartments, where they fold, this may reflect the cytosolic and mitochondrial fractions of mitochondrial proteins. In classical LiP experiments performed in lysates, both fractions should be equally accessible to PK. In the in-cell context however, only the cytosolic and nuclear fractions may be accessible, in particular since the double mitochondrial membrane may limit PK uptake upon electroporation. Consistent with this, there was no enrichment of the term “mitochondria” within PK-cleaved proteins in the in-cell LiP setup (Fig 1f).

Next, we asked if LiP peptides reflective of low structural similarity between in-cell LiP and classical LiP were enriched in specific domains. We excluded mitochondrial proteins from this analysis because we cannot distinguish functional differences from differences in localization. Domain-level gene ontology enrichment analysis for peptides with a higher intensity in in-cell LiP (ratio > 1.1 of log_10_-transformed peptide intensities) showed a significant enrichment for RNA and DNA binding domains (Figure 3d). This indicates that in-cell LiP provides information on RNA and DNA interactions with proteins, which are likely affected in classical LiP experiments due to cells lysis and centrifugation.

### In-cell LiP-MS identifies structural changes in stress granules

Our data on the rapamycin-FKB1A interaction showed that the in-cell LIP approach can detect structural alterations in proteins within the cell induced by a high-affinity interaction, which is also detectable after cell lysis. The promise of an in-cell method for detecting structural alterations is that it may also be able to capture the dynamics of biomolecular condensates that are too complex to properly reconstitute in vitro, and that may be partly lost upon cell lysis (Jain et al., 2016).We therefore asked whether in-cell LiP-MS can detect structural changes of proteins that interact in stress granules. Stress granules are cytosolic biomolecular condensates composed of proteins and RNA that can be observed in response to various stresses. Their formation is probably at least partly driven by liquid-liquid phase separation via low affinity interactions (Glauninger et al., 2022). We treated HEK293 cells with sodium arsenite, which is known to induce the formation of stress granules, and incubated them under these stress conditions for up to 90 minutes. At different time points (0, 10, 20 and 90 minutes after arsenite treatment), cells were detached in PBS pre-warmed to 37 °C, centrifuged at 200 x g for 30 seconds, resuspended in PBS, immediately electroporated with PK and then processed by in-cell LiP. We confirmed that stress granules formed between 10 and 20 minutes of arsenite treatment using immunofluorescence staining for the stress granule marker G3BP1 (Figure 4a), which is in line with previous observations (Jain et al., 2016).

**Figure 4:**
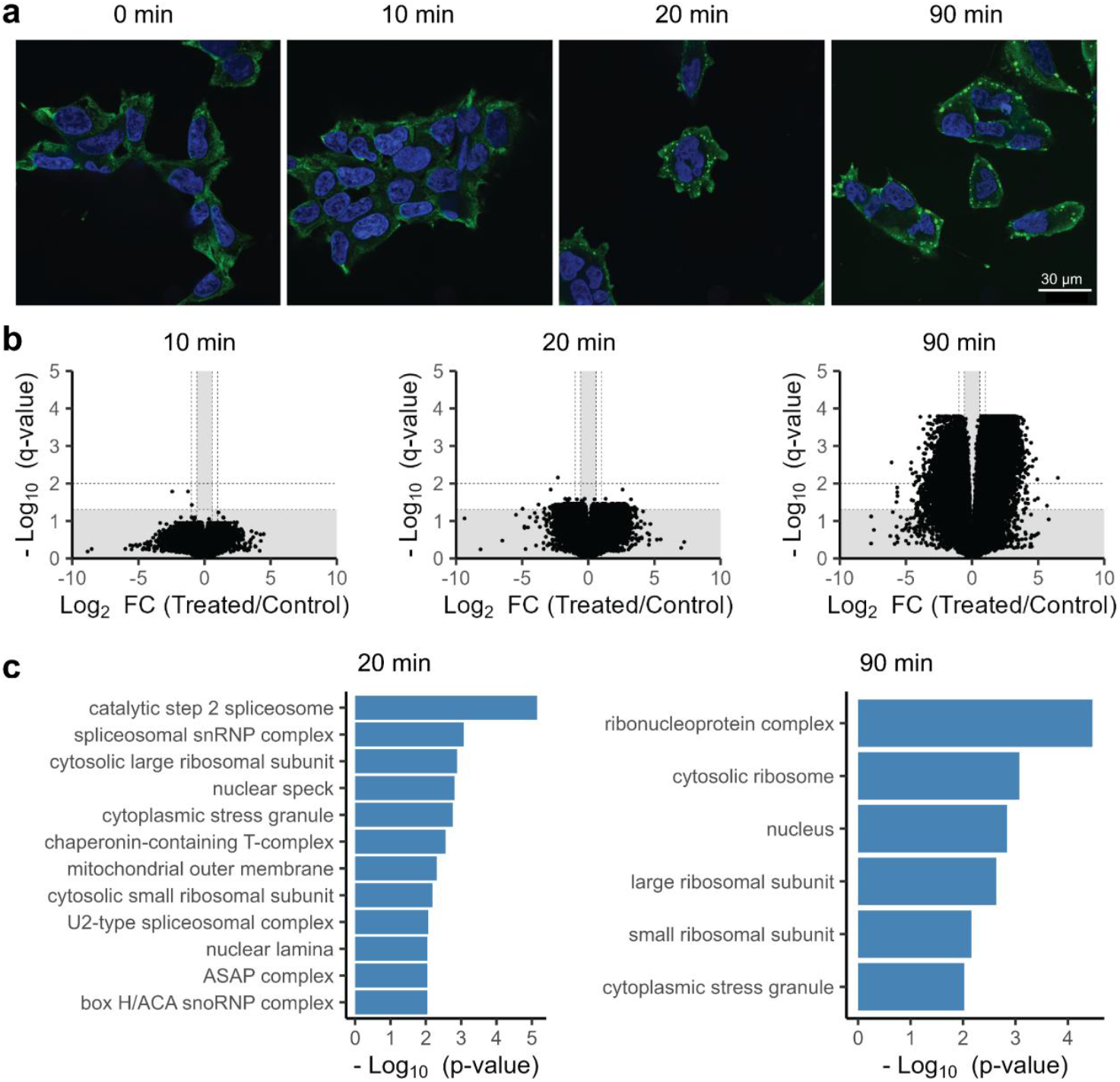
In-cell LiP-MS analysis of structural changes upon arsenite treatment of mammalian cells. (**a**) Representative micrographs of HEK293 cells stained with anti-G3BP1 antibody (green) and Hoechst (blue) at the indicated time points of sodium arsenite treatment. Scale bar, 30 µm. **(b)** Peptide intensities from HEK293 cells treated with sodium arsenite compared to untreated cells at 10, 20, and 90 min. Each data point represents a single peptide. The shaded gray region marks significance levels (FC ≥ 1.5, p-value < 0.05, n = 6 replicates). **(c)** Gene ontology enrichment analysis (cellular component) of proteins showing structural changes upon arsenite treatment. (p-value < 0.01).

After 10 minutes of treatment, before stress granules were observed by microscopy, we detected structural changes in only three peptides from three proteins in arsenite-treated samples relative to untreated control; these were Creatine kinase B-type (CKB), Protein phosphatase 1 regulatory subunit 12A (PPP1R12A), and SERPINE1 mRNA-binding protein 1 (SERBP1) (Figure 4b; FC>1.5, q-value < 0.05). At 20 minutes of arsenite treatment, 2325 peptides on 1149 proteins had changed significantly, rising to 20,841 peptides on 3146 proteins at 90 minutes of treatment. With few exceptions (1 of 4204 proteins at 10 min, 13 of 4207 proteins at 90 min arsenite treatment), protein levels did not change significantly in this time frame (Figure S6a; FC > 1.5, q-value < 0.05, n = 6 replicates), and we therefore did not globally normalize protein structural changes for protein abundance changes.

Gene ontology enrichment analysis showed that proteins that had changed structure after 20 minutes and 90 minutes of arsenite treatment were indeed enriched in components of stress granules (Figure 4c). Moreover, proteins involved in RNA binding and RNA-related biological processes, again associated with stress granules, were also significantly enriched (Figure S6b, c). We observed that more cleavage events were detected at later time points as measured by the fraction of HT peptides (average percentage of HT peptides was 1.7-fold higher at 90 min arsenite treatment than in untreated samples; Figure S6d), suggesting that the proteome of HEK cells may become slightly more susceptible to PK cleavage upon arsenite treatment.

Next, we asked whether in-cell LiP identified structural changes in known stress granule-associated proteins. Interestingly, we captured changes consistent with the known mechanism of G3BP1 (Figure 5a), a core component of stress granule assembly. Stress granule formation is triggered by a structural change in G3BP1 in response to free RNA. G3BP1 consists of a NTF2-like domain, which is required for dimerization (Figure 5b), an RNA recognition motif, and three intrinsically disordered regions (IDRs). Both dimerization and RNA binding by G3BP1 are essential for stress granule assembly (Sanders et al., 2020). It has been suggested that G3BP1 adopts a closed conformation in the absence of stress, held together by electrostatic interactions between IDR1 and IDR3. When free RNA levels increase during arsenite stress, the interaction between the IDRs is disrupted and RNA binds to the RNA recognition motif and. The RNA-bound open state of G3BP1 serves as the core of the stress granule and leads to condensation (Yang et al., 2020). After 20 minutes of arsenite treatment, we detected only one altered LiP peptide on G3BP1: this peptide is located exactly on the RNA recognition motif of the protein (Figure 5c), in line with this proposed model of structural rearrangements making the RNA binding domain accessible upon stress. At 90 minutes, structural changes occurred within all domains of G3BP1 (Figure 5c). This suggests that structural changes associated with biomolecular condensation can be detected by in-cell LiP-MS.

**Figure 5:**
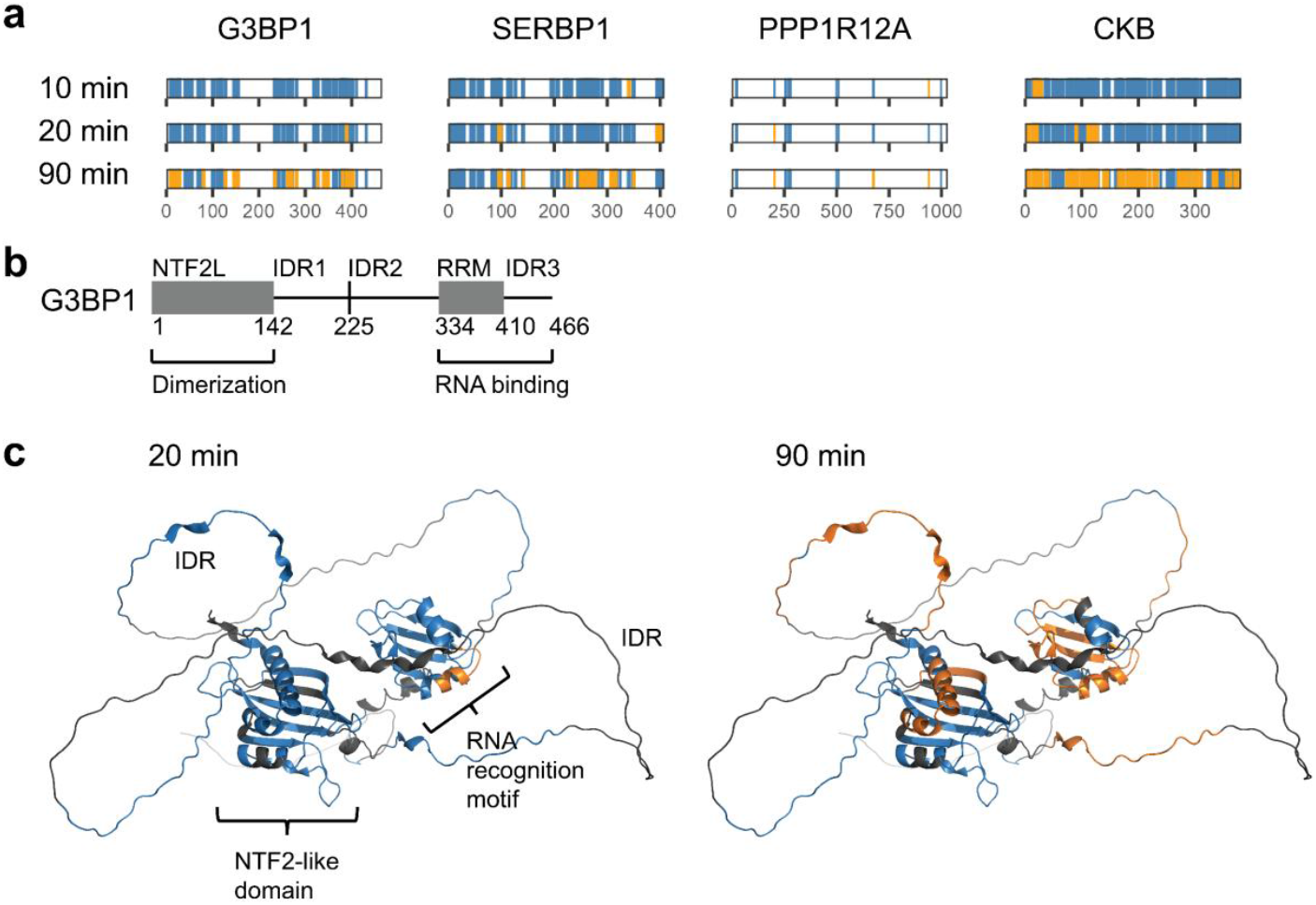
Structural changes upon sodium arsenite treatment of HEK293 cells. (**a**) Barcode plots of protein structural changes upon arsenite treatment. The blue regions indicate peptides detected by mass spectrometry. Orange regions changed significantly at the indicated time (FC ≥ 1.5, p-value < 0.05, n = 6 replicates). (**b**) Domains of G3BP1. NTF2L: NTF2-like domain; IDR: intrinsically disordered region; RRM: RNA recognition motif. (**c**) Alphafold structure of G3BP1. Blue regions were detected by mass spectrometry. Orange regions changed significantly at the indicated time.

More broadly, in-cell LiP-MS identified novel structural changes in 190 of 200 high confidence stress granule-associated proteins (based on the RNAgranuleDB v2.0 data base) that are also MS-detectable (Millar et al., 2023). We obtained peptide-level resolution of protein structure dynamics, as shown for 35 stress granule proteins selected manually for their special interest (Figure S7). Although stress granules are not detected by microscopy until after 20 minutes of arsenite treatment, significantly changed LiP peptides were detected on three proteins, SERBP1, PPP1R12A, and CKB, after 10 minutes (Figure 5a). PPP1R12A associates with stress granules (Yang et al., 2020). SERBP1 acts as a ribosome dormancy or preservation factor upon mTOR phosphorylation (Shetty et al., 2023) and, upon arsenite treatment, localizes to both stress granules and nucleoli (Lee et al., 2014). SERBP1 has also been implicated in clearance of stress granules after arsenite stress by modulating the degradation of G3BP1 (Wang et al., 2023). To our knowledge, CKB has not been previously associated with stress granules. Thus, we provide information on the structure dynamics of novel and known stress granule proteins during arsenite stress, with many changes mapping to known disordered and RNA binding regions (Figure 6).

**Figure 6:**
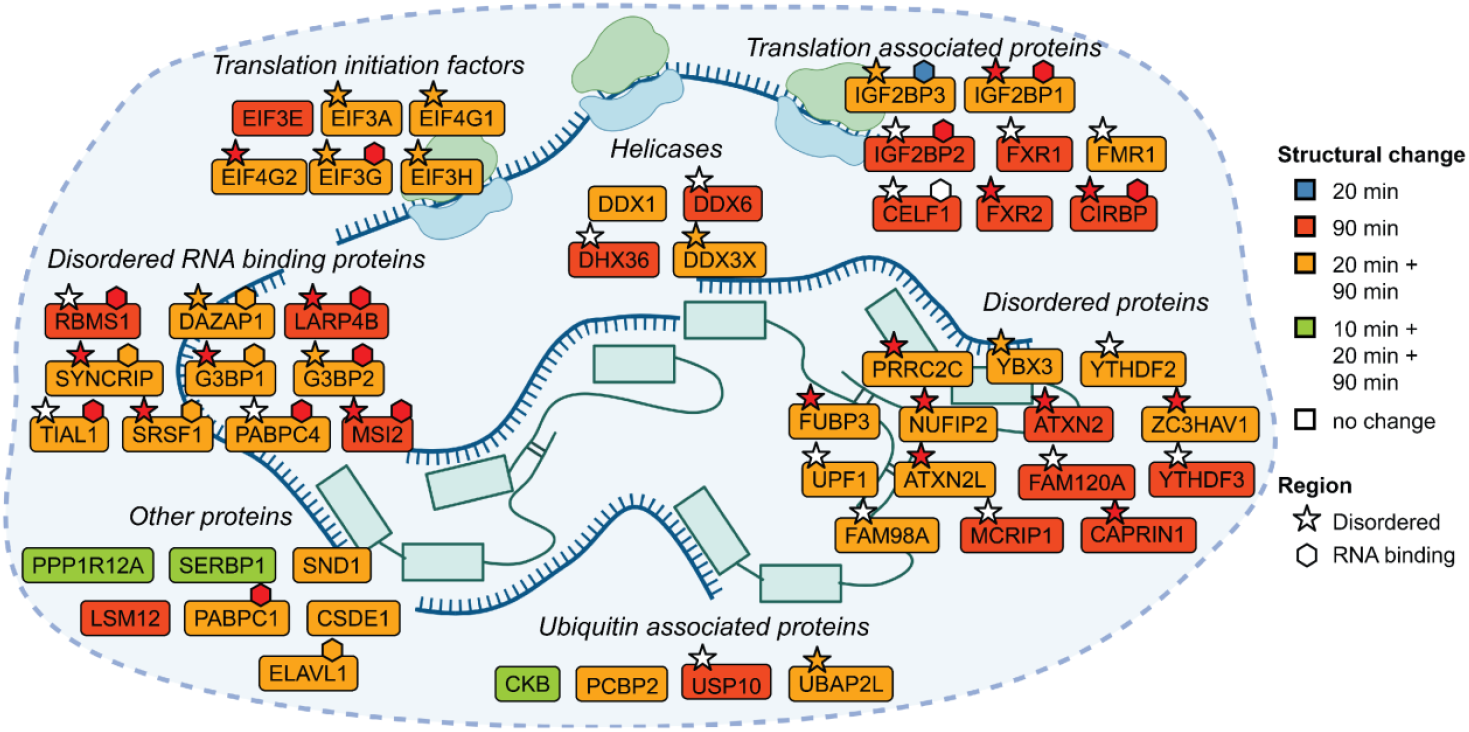
Structural changes in stress granule proteins. Illustration of a stress granule at different times of sodium arsenite treatment. A star shape indicates at least one disordered region annotated in Uniprot, a hexagon at least one RNA binding region annotated in Uniprot. Colored proteins change significantly in structure at the indicated time (FC ≥ 1.5, p-value < 0.05, n = 6 replicates; blue: changes only at 20 min; red changes only at 90 min; green: changes at 10 min, 20 min, and 90 min; orange: changes at 20 min and 90 min). Colored stars and hexagons indicate that at least one significantly changing peptide is located in the indicated region. Depicted are proteins that are reported to be associated in RNAgranuleDB v2.0 with BioID or microscopy evidence on this website RNAgranuleDB v2.0 was further filtered for proteins that were detected in this study, and the fifty proteins with the highest score on RNAgranuleDB 2.0 were selected. Additionally, CKB, PPP1R12A, and SERBP1 are shown.

Interestingly, gene ontology analysis also suggested that nuclear speckles, another membraneless biomolecular condensate, and the spliceosome, which associates with nuclear speckles, are enriched among proteins that show structural changes after 20 minutes of arsenite treatment. At this time point, we observed changes in 80 proteins that are associated with nuclear speckles (based on gene ontology annotations), including 14 splicing factors, and the phosphatase PP1-gamma catalytic subunit, which has been reported to dynamically re-localize between cytoplasm and nuclear speckles (Trinkle-Mulcahy et al., 2001). After 90 min, 161 proteins located in nuclear speckles showed evidence of structural changes, including the known marker protein SC35. To assess whether these in-cell LiP detected structural alterations of nuclear speckle proteins detected by in-cell LiP reflected morphological changes in this phase-separated compartment, we monitored arsenite-treated cells with fluorescence microscopy after staining for the known nuclear speckle marker SC35 (Figure 7a). We assessed the circularity of nuclear speckles, which is known to change upon alterations in the dynamics of this compartment. We found that nuclear speckles became rounder upon sodium arsenite treatment in comparison to untreated control cells (Figure 7a, b). This effect is in line with reported changes of nuclear speckle morphology upon other stresses (J. Kim et al., 2019; Raina & Rao, 2022; Spector et al., 1991). Taken together, our data show that in-cell LiP identified both known and novel alterations in proteins connected to biomolecular condensates within human cells. Further, we have used this approach to generate a compendium of proteins undergoing structural alterations upon arsenite stress, and to identify structurally altered regions in these proteins.

**Figure 7:**
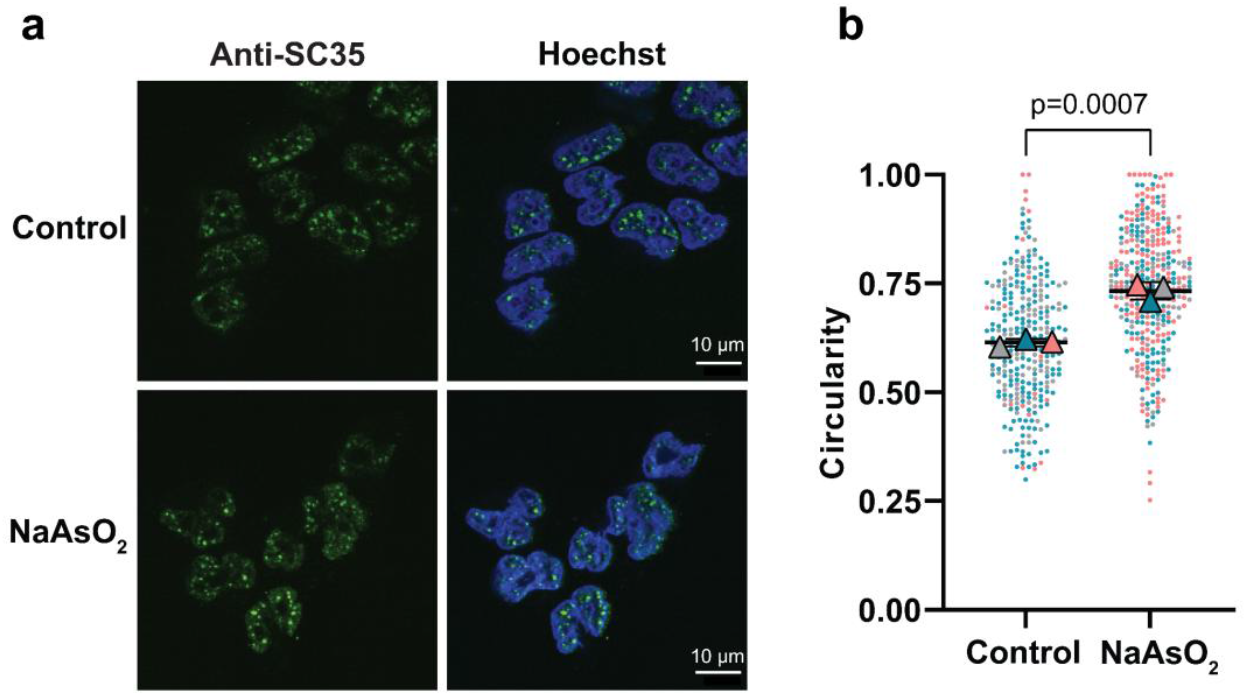
Nuclear speckles become rounder upon sodium arsenite treatment. (**a**) Representative images of HEK293 cells stained with anti-SC35 antibody and Hoechst after 90 min treatment with sodium arsenite. Scale bars, 10 µm. (**b**) Circularity of nuclear speckles with and without sodium arsenite treatment measured for three biological replicates. Colors correspond to replicates. Two-tailed t-test was performed with the means of each replicate. Error bars indicate the standard deviation (*** p-value = 0.0007).

## Discussion

We present in-cell LiP-MS, a novel method to study proteome-scale structural changes in their native environment. Compared to LiP-MS in lysates, intracellular organization and molecular concentrations are preserved in this approach, enabling the study of changes in cellular physiology with minimal invasion. Since LiP-MS captures functional molecular events such as enzyme activity, protein-protein interaction, PTM and allosteric regulation, as we have previously shown (Cappelletti et al., 2021), the in-cell version of this technique should prove a powerful method for various types of functional proteomics studies.

Although some existing structural proteomic methods can be implemented in intact cells, they face challenges. Hydroxyl radical footprinting (HRF)-MS reports on intracellular structural changes with residue-level resolution, but requires a microfluidic setup and oxidations are only seen in approximately 100 proteins (Espino et al., 2015). Crosslinking mass spectrometry applied to intact cells also suffers from low proteome coverage. Thermal protein profiling (TPP) uses a simple temperature gradient to identify proteins that change stability under conditions of interest and does not suffer from low coverage, but it lacks structural information since it does not pinpoint the region(s) of a protein that may be altered (Savitski et al., 2014). Further, our approach has advantages over in-cell NMR, since in-cell LiP is applied on proteome-scale and does not require labeling. However, the two approaches may be complementary since in-cell LiP does not achieve the atomic resolution of NMR. Structurally altered proteins that are detected by in-cell LiP in an unbiased manner can be followed up by in-cell NMR.

We expect that modification of several parameters could help extend our approach in the future. With the current settings, we identified several thousands of proteins, however, sequence coverage per protein remained medium to low in most cases. While coverage was not increased with higher PK concentration and prolonged incubation time, in applications where cells do not need to remain viable after electroporation, higher electroporation voltage and pulse number may lead to more uniform influx of PK and possibly to better sequence coverage.

In-cell LiP-MS may be adapted to other temperatures, pH, or different low-specificity proteases. It may even be feasible to introduce an inactivated protease by electroporation, such that the start of proteolysis can be more tightly controlled. For example, the catalytic residues of a protease may be replaced with a photocaged homologue such that it can be light-activated under conditions of interest. Further, in-cell LiP could be used not only to cell lines, and may be applicable also to tissues and to unicellular organisms such as bacteria or yeast. The FRET-based proteolysis assay we have described here should prove useful to screen for optimal conditions to adapt in-cell LiP to other systems. The reporter may also be used to test whether experimental conditions affect PK activity which, as in classical LiP-MS, can affect the interpretation of an experiment.

We have optimized in-cell LiP-MS conditions to preserve the native state of cells as far as possible. For example, care should be taken that cells do not clump after incubation with extracellular PK, most probably due to partial proteolysis of surface proteins. Clumping during limited proteolysis can be avoided if the added PK is ice-cold, electroporation is performed within seconds after mixing PK and cells, and cells are resuspended well prior to addition of PK. Good cell resuspension also results in higher PK uptake. Mixing after protease addition should be performed within seconds and further mixing should be avoided.

In classical LiP-MS on lysates, optimal coverage has been found for 1 μg PK added to 100 μg substrate protein. For in-cell LiP, we obtained the highest coverage by electroporating 3 million HEK293 cells with 12 nmol PK, of which 36.6 pmol PK were intracellular. If different cell types or electroporation settings are used, we recommend first screening for conditions that yield good PK internalization using the FRET reporter, and then optimizing the conditions using the % half-tryptic peptides or the number of PK cleavages in the proteome as a readout. Coverage can then be determined using mass spectrometry by comparing cells electroporated with PK to control cells to which PK was added but not electroporated. Another difference between in-cell and in-lysate LiP is the homogeneity of the system: in lysates, an almost homogeneous mixture of proteins, PK and any other added perturbant can be achieved, whereas in-cell LiP analyzes a mixture of cells which may be in different cellular states, and PK uptake may vary. Perhaps as a result, reproducibility is currently higher in classical LiP, which in part can be compensated by using more replicates in in-cell LiP. Because exact timing matters in LiP, up to eight lysates were treated and processed simultaneously. In our current setup, the electroporation equipment limits us to electroporating one sample at a time and thus we anticipate that increasing the throughput of this step would improve reproducibility.

Classical LiP-MS can be used to identify drug targets in cell lysates. The in-cell LiP approach detected the binding of rapamycin to FKBP1A, its only known target in mammalian cells, showing that the method detects a conformational change on a single target protein in a very complex and crowded background. While the effect was smaller than seen for LiP-MS in lysates, this could be because of sample heterogeneity or because protease accessibility of FKBP1A may be different in the crowded environment of the cell. Moreover, the intracellular target might be less saturated than in the lysate where the drug is added in excess to a homogeneous protein suspension.

In-cell LiP-MS should have the ability to identify not only direct binding partners but also downstream effects, which may be enhanced and separated from target-engagement effects with time-course experiments. This opens the exciting possibility that the approach could be used to identify drug targets or targeted pathways, downstream events, and off-target effects within intact cells under specific conditions of interest, e.g., hypoxia. We have previously shown that dose-response experiments can help rank potential binding partners in classical LiP-MS, and we envision that such a strategy could be applied in the in-cell context as well. Dose- and time-resolved drug profiling of post-translational modifications (PTMs) has been used to group proteins involved in different downstream events (Zecha et al., 2023). This approach allowed to elucidate the mode of action of drugs in cells and even to assign PTMs to pathways.

For drugs that trigger fast intracellular responses, in-cell LiP analyses may result in large sets of hits even at early time points, which could preclude the separation of direct binding and downstream events. In such cases, target engagement events can be prioritized by LiP-MS experiments performed in parallel on drug-treated cell lysates. In-cell LiP-MS could also help to elucidate effects of mutations in pathways affected by drug binding, as well as the effect of two or more drugs added simultaneously or subsequently to cells.

The study also generated a resource that pinpoints proteins whose structure is affected if they are studied outside the cellular environment, and on RNA and DNA interacting domains that are altered upon cell lysis. Developing organelle-specific in cell LiP by targeting the protease to the different intracellular locations could be an interesting direction to detect organelle-specific structural states and in general to probe protein structures in subcellular compartments that are currently not accessible to our setup.

Application of in-cell LiP-MS to arsenite-stressed cells provides to our knowledge the first *in situ* insight into structural rearrangements of stress granule proteins. These dynamic structures have typically been studied by fluorescence microscopy, which lacks structural resolution, or by *in vitro* reconstitution, but until now there were no methods available to study structure information in large dynamic assemblies. Stress granule proteins, including the scaffolding protein G3BP1, were enriched in the set of structurally altered proteins upon arsenite stress, suggesting that the dataset may be used to generate hypotheses about so far unknown components or regulators of stress granules. Information on the specific protein regions involved may shed light on mechanisms of stress granule formation. Overall, these data provide the first resource on proteome-scale and time-resolved structural changes in human cells under stress with peptide-level resolution.

Of special interest are the structural changes in CKB, PPP1R12A and SERBP1, detected before stress granules are visible by fluorescence microscopy. The enzyme Creatine kinase B-type (CKB) plays an important role in energy transfer and homeostasis and has not so far been linked to stress granules. CKB transfers phosphate from ATP to creatine. The resulting phosphocreatine serves as donor of the phosphate group to rapidly regenerate ATP from ADP. Although the observed changes could be in principle caused by sodium arsenite unrelated to stress granules, the timing suggests that the protein might be involved in the stress granule formation. The Protein phosphatase 1 regulatory subunit 12A (PPP1R12A) is involved in various cellular processes including muscle contraction and cell cycle regulation, where it antagonizes Polo-like kinase 1 (PLK1), an essential regulator of mitosis (Yamashiro et al., 2008). PPP1R12A has been detected as part of stress granules (Yang et al., 2020) but its functional role is unknown; our data suggest that it may play a role in the early stages of stress granule formation and pinpoints the regions involved. The SERPINE1 mRNA-binding protein 1 (SERBP1), is known to interact with stress-granule proteins upon arsenite treatment in an RNA-dependent manner (Lee et al., 2014) and has been described to be essential for stress granule clearance after arsenite stress (Wang et al., 2023). The same study found that knocking out SERBP1 had no effect on stress granules after 60 min of arsenite treatment, consistent with our observation of an earlier structural change in this protein. It is also possible that the structural change we detect in SERBP1 occurs in the nuclear-located protein, which is known to increase in level upon arsenite stress, and where it likely it binds to mRNA free ribosomes to stabilize the 80S particle (Shetty et al., 2023), or due to the relocation itself (Lee et al., 2014).

Nuclear speckles are dynamic assemblies of splicing factors that were not yet implicated in the response to arsenite stress, but we observed enrichment of both speckle proteins and the splicing machinery among structurally altered proteins upon arsenite treatment. In line with this, we could observe morphological changes in these compartments with fluorescence microscopy, which have previously been seen upon heat shock (Spector et al., 1991), upon inhibition of transcription (T. Kim et al., 2023), and upon inhibition of the DEAD-box ATPase UAP56 (Hondele et al., 2019). Further analysis of the set of structurally altered proteins may provide insight into mechanisms leading to the altered morphology of nuclear speckles and the molecular players involved.

In the future we envision a broad range of applications of in-cell LiP in the field of condensates. For example, our data may be used to identify LiP marker peptides for stress granule formation, potentially including markers that probe the involvement of the RNA binding and IDP regions for each protein. Similar markers could be identified for the formation of other physiological molecular condensates (such as Cajal bodies or p-bodies) or metabolons (e.g. the purinosome) and even for pathological condensates (e.g. from amyloidogenic proteins such as FUS or TDP43). These markers, potentially measured simultaneously with a targeted MS analysis, could be used to study the detailed dynamics of such structures within the cell, identify perturbations that influence them, and study potential cross-talk between condensates. One may envision using such markers to derive in cell phase diagrams of condensates.

Other applications of the technique beyond condensates may include the characterization of the global cellular response to changes in pH, such as occurs in the tumor microenvironment, or to other environmental factors such as toxins or irradiation. In-cell LiP may provide mechanistic information into viral entry, receptor internalization upon drug binding, the binding of modified T-cells in immunotherapy, or nutrient shifts. Another exciting application of the approach is the study of altered material properties of the cytosol, e.g. the phenomenon of extreme cytosol rigidification upon starvation and quiescence (Joyner et al., 2016; Munder et al., 2016). The nature of protein structures in this state is still unknown; they may phase separate or aggregate, this may involve all proteins or only a subset, and RNA-binding, intrinsically disordered or other protein domains could be involved. In-cell LiP-MS detected structural changes could shed light on all of these questions.

In summary, we have shown that in-cell LiP-MS can detect proteome-wide structural changes upon a perturbation of interest, within the native environment of the intact cell and at peptide resolution. Since protein structural changes are likely to be ubiquitous under perturbation or stimulus of a biological system, in-cell LiP-MS has the potential to provide insight into numerous cellular processes including labile assemblies formed by phase separation that could so far not be addressed with established methods.

## Acknowledgements

We thank Walther Hänseler (University of Zurich) for performing initial electroporation experiments, Juan Gerez (ETH Zurich) for advice on electroporation settings, Roland Riek (ETH Zurich) for providing access to the electroporation equipment, Donald Hilvert (ETH Zurich) for insightful discussions, Charlie Morgan (Australian National University) and Jason Chin (MRC-LMB Cambridge) for sharing the ECFP-YPet plasmid for sensing protease activity, and Federico Uliana (ETH Zurich) for advice on the absolute quantification of proteinase K.

This work was funded by the European Research Council (866004), and the EPIC-XS Consortium (823839), the last two under the EU Horizon 2020 program, and the National Center of Competence in Research AntiResist funded by the SNSF (grant numbers 51NF40_180541 and TMAG-3_209354).

## References

Baek, M., DiMaio, F., Anishchenko, I., Dauparas, J., Ovchinnikov, S., Lee, G. R., Wang, J., Cong, Q., Kinch, L. N., Schaeffer, R. D., Millán, C., Park, H., Adams, C., Glassman, C. R., DeGiovanni, A., Pereira, J. H., Rodrigues, A. V., van Dijk, A. A., Ebrecht, A. C., … Baker, D. (2021). Accurate prediction of protein structures and interactions using a three-track neural network. Science, 373(6557), 871–876. 10.1126/science.abj8754

Cappelletti, V., Hauser, T., Piazza, I., Pepelnjak, M., Malinovska, L., Fuhrer, T., Li, Y., Dörig, C., Boersema, P., Gillet, L., Grossbach, J., Dugourd, A., Saez-Rodriguez, J., Beyer, A., Zamboni, N., Caflisch, A., de Souza, N., & Picotti, P. (2021). Dynamic 3D proteomes reveal protein functional alterations at high resolution in situ. Cell, 184(2), 545-559.e22. 10.1016/j.cell.2020.12.021

Espino, J. A., Mali, V. S., & Jones, L. M. (2015). In Cell Footprinting Coupled with Mass Spectrometry for the Structural Analysis of Proteins in Live Cells. Analytical Chemistry, 87(15), 7971–7978. 10.1021/acs.analchem.5b01888

Feng, Y., De Franceschi, G., Kahraman, A., Soste, M., Melnik, A., Boersema, P. J., de Laureto, P. P., Nikolaev, Y., Oliveira, A. P., & Picotti, P. (2014). Global analysis of protein structural changes in complex proteomes. Nature Biotechnology, 32(10), 1036–1044. 10.1038/nbt.2999

Glauninger, H., Wong Hickernell, C. J., Bard, J. A. M., & Drummond, D. A. (2022). Stressful steps: Progress and challenges in understanding stress-induced mRNA condensation and accumulation in stress granules. Molecular Cell, 82(14), 2544–2556. 10.1016/j.molcel.2022.05.014

Ha, T., Ting, A. Y., Liang, J., Caldwell, W. B., Deniz, A. A., Chemla, D. S., Schultz, P. G., & Weiss, S. (1999). Single-molecule fluorescence spectroscopy of enzyme conformational dynamics and cleavage mechanism. Proceedings of the National Academy of Sciences, 96(3), 893–898. 10.1073/pnas.96.3.893

Heitman, J., Movva, N. R., & Hall, M. N. (1991). Targets for Cell Cycle Arrest by the Immunosuppressant Rapamycin in Yeast. Science, 253(5022), 905–909. 10.1126/science.1715094

Hondele, M., Sachdev, R., Heinrich, S., Wang, J., Vallotton, P., Fontoura, B. M. A., & Weis, K. (2019). DEAD-box ATPases are global regulators of phase-separated organelles. Nature, 573(7772), 144–148. 10.1038/s41586-019-1502-y

Jain, S., Wheeler, J. R., Walters, R. W., Agrawal, A., Barsic, A., & Parker, R. (2016). ATPase-Modulated Stress Granules Contain a Diverse Proteome and Substructure. Cell, 164(3), 487–498. 10.1016/j.cell.2015.12.038

Joyner, R. P., Tang, J. H., Helenius, J., Dultz, E., Brune, C., Holt, L. J., Huet, S., Müller, D. J., & Weis, K. (2016). A glucose-starvation response regulates the diffusion of macromolecules. ELife, 5(MARCH2016). 10.7554/eLife.09376

Kim, J., Han, K. Y., Khanna, N., Ha, T., & Belmont, A. S. (2019). Nuclear speckle fusion via long-range directional motion regulates speckle morphology after transcriptional inhibition. Journal of Cell Science, 132(8). 10.1242/jcs.226563

Kim, T., Yoo, J., Do, S., Hwang, D. S., Park, Y., & Shin, Y. (2023). RNA-mediated demixing transition of low-density condensates. Nature Communications, 14(1), 2425. 10.1038/s41467-023-38118-z

Lee, Y., Wei, H., Chen, L., & Li, C. (2014). Localization of SERBP1 in stress granules and nucleoli. The FEBS Journal, 281(1), 352–364. 10.1111/febs.12606

Mackmull, M.-T., Nagel, L., Sesterhenn, F., Muntel, J., Grossbach, J., Stalder, P., Bruderer, R., Reiter, L., van de Berg, W. D. J. J., de Souza, N., Beyer, A., & Picotti, P. (2022). Global, in situ analysis of the structural proteome in individuals with Parkinson’s disease to identify a new class of biomarker. Nature Structural & Molecular Biology, 29(10), 978–989. 10.1038/s41594-022-00837-0

Malinovska, L., Cappelletti, V., Kohler, D., Piazza, I., Tsai, T.-H., Pepelnjak, M., Stalder, P., Dörig, C., Sesterhenn, F., Elsässer, F., Kralickova, L., Beaton, N., Reiter, L., de Souza, N., Vitek, O., & Picotti, P. (2023). Proteome-wide structural changes measured with limited proteolysis-mass spectrometry: an advanced protocol for high-throughput applications. Nature Protocols, 18(3), 659–682. 10.1038/s41596-022-00771-x

Millar, S. R., Huang, J. Q., Schreiber, K. J., Tsai, Y.-C., Won, J., Zhang, J., Moses, A. M., & Youn, J.-Y. (2023). A New Phase of Networking: The Molecular Composition and Regulatory Dynamics of Mammalian Stress Granules. Chemical Reviews, 123(14), 9036–9064. 10.1021/acs.chemrev.2c00608

Munder, M. C., Midtvedt, D., Franzmann, T., Nüske, E., Otto, O., Herbig, M., Ulbricht, E., Müller, P., Taubenberger, A., Maharana, S., Malinovska, L., Richter, D., Guck, J., Zaburdaev, V., & Alberti, S. (2016). A pH-driven transition of the cytoplasm from a fluid-to a solid-like state promotes entry into dormancy. ELife, 5(MARCH2016). 10.7554/eLife.09347

Piazza, I., Beaton, N., Bruderer, R., Knobloch, T., Barbisan, C., Chandat, L., Sudau, A., Siepe, I., Rinner, O., de Souza, N., Picotti, P., & Reiter, L. (2020). A machine learning-based chemoproteomic approach to identify drug targets and binding sites in complex proteomes. Nature Communications, 11(1), 4200. 10.1038/s41467-020-18071-x

Piazza, I., Kochanowski, K., Cappelletti, V., Fuhrer, T., Noor, E., Sauer, U., & Picotti, P. (2018). A Map of Protein-Metabolite Interactions Reveals Principles of Chemical Communication. Cell, 172(1–2), 358-372.e23. 10.1016/j.cell.2017.12.006

Raina, K., & Rao, B. J. (2022). Mammalian nuclear speckles exhibit stable association with chromatin: a biochemical study. Nucleus, 13(1), 58–73. 10.1080/19491034.2021.2024948

Sakakibara, D., Sasaki, A., Ikeya, T., Hamatsu, J., Hanashima, T., Mishima, M., Yoshimasu, M., Hayashi, N., Mikawa, T., Wälchli, M., Smith, B. O., Shirakawa, M., Güntert, P., & Ito, Y. (2009). Protein structure determination in living cells by in-cell NMR spectroscopy. Nature, 458(7234), 102–105. 10.1038/nature07814

Sanders, D. W., Kedersha, N., Lee, D. S. W., Strom, A. R., Drake, V., Riback, J. A., Bracha, D., Eeftens, J. M., Iwanicki, A., Wang, A., Wei, M.-T., Whitney, G., Lyons, S. M., Anderson, P., Jacobs, W. M., Ivanov, P., & Brangwynne, C. P. (2020). Competing Protein-RNA Interaction Networks Control Multiphase Intracellular Organization. Cell, 181(2), 306-324.e28. 10.1016/j.cell.2020.03.050

Savitski, M. M., Reinhard, F. B. M., Franken, H., Werner, T., Savitski, M. F., Eberhard, D., Molina, D. M., Jafari, R., Dovega, R. B., Klaeger, S., Kuster, B., Nordlund, P., Bantscheff, M., & Drewes, G. (2014). Tracking cancer drugs in living cells by thermal profiling of the proteome. Science, 346(6205). 10.1126/science.1255784

Schopper, S., Kahraman, A., Leuenberger, P., Feng, Y., Piazza, I., Müller, O., Boersema, P. J., & Picotti, P. (2017). Measuring protein structural changes on a proteome-wide scale using limited proteolysis-coupled mass spectrometry. Nature Protocols, 12(11), 2391–2410. 10.1038/nprot.2017.100

Shetty, S., Hofstetter, J., Battaglioni, S., Ritz, D., & Hall, M. N. (2023). TORC1 phosphorylates and inhibits the ribosome preservation factor Stm1 to activate dormant ribosomes. The EMBO Journal, 42(5), e112344. 10.15252/embj.2022112344

Spector, D. L., Fu, X. D., & Maniatis, T. (1991). Associations between distinct pre-mRNA splicing components and the cell nucleus. The EMBO Journal, 10(11), 3467–3481. 10.1002/j.1460-2075.1991.tb04911.x

Trinkle-Mulcahy, L., Sleeman, J. E., & Lamond, A. I. (2001). Dynamic targeting of protein phosphatase 1 within the nuclei of living mammalian cells. Journal of Cell Science, 114(23), 4219–4228. 10.1242/jcs.114.23.4219

Tunyasuvunakool, K., Adler, J., Wu, Z., Green, T., Zielinski, M., Žídek, A., Bridgland, A., Cowie, A., Meyer, C., Laydon, A., Velankar, S., Kleywegt, G. J., Bateman, A., Evans, R., Pritzel, A., Figurnov, M., Ronneberger, O., Bates, R., Kohl, S. A. A., … Hassabis, D. (2021). Highly accurate protein structure prediction for the human proteome. Nature, 596(7873), 590–596. 10.1038/s41586-021-03828-1

Wang, F., Wang, L., Gan, S., Feng, S., Ouyang, S., Wang, X., & Yuan, S. (2023). SERBP1 Promotes Stress Granule Clearance by Regulating 26S Proteasome Activity and G3BP1 Ubiquitination and Protects Male Germ Cells from Thermostimuli Damage. Research, 6, 1–18. 10.34133/research.0091

Yamashiro, S., Yamakita, Y., Totsukawa, G., Goto, H., Kaibuchi, K., Ito, M., Hartshorne, D. J., & Matsumura, F. (2008). Myosin Phosphatase-Targeting Subunit 1 Regulates Mitosis by Antagonizing Polo-like Kinase 1. Developmental Cell, 14(5), 787–797. 10.1016/j.devcel.2008.02.013

Yang, P., Mathieu, C., Kolaitis, R.-M., Zhang, P., Messing, J., Yurtsever, U., Yang, Z., Wu, J., Li, Y., Pan, Q., Yu, J., Martin, E. W., Mittag, T., Kim, H. J., & Taylor, J. P. (2020). G3BP1 Is a Tunable Switch that Triggers Phase Separation to Assemble Stress Granules. Cell, 181(2), 325-345.e28. 10.1016/j.cell.2020.03.046

Zecha, J., Bayer, F. P., Wiechmann, S., Woortman, J., Berner, N., Müller, J., Schneider, A., Kramer, K., Abril-Gil, M., Hopf, T., Reichart, L., Chen, L., Hansen, F. M., Lechner, S., Samaras, P., Eckert, S., Lautenbacher, L., Reinecke, M., Hamood, F., … Kuster, B. (2023). Decrypting drug actions and protein modifications by dose- and time-resolved proteomics. Science, 380(6640), 93–101. 10.1126/science.ade3925

