## Supplemental Figures and Methods for "Limited proteolysis-coupled mass spectrometry captures proteome-wide protein structural alterations and biomolecular condensation in living cells"

### Supplementary figures

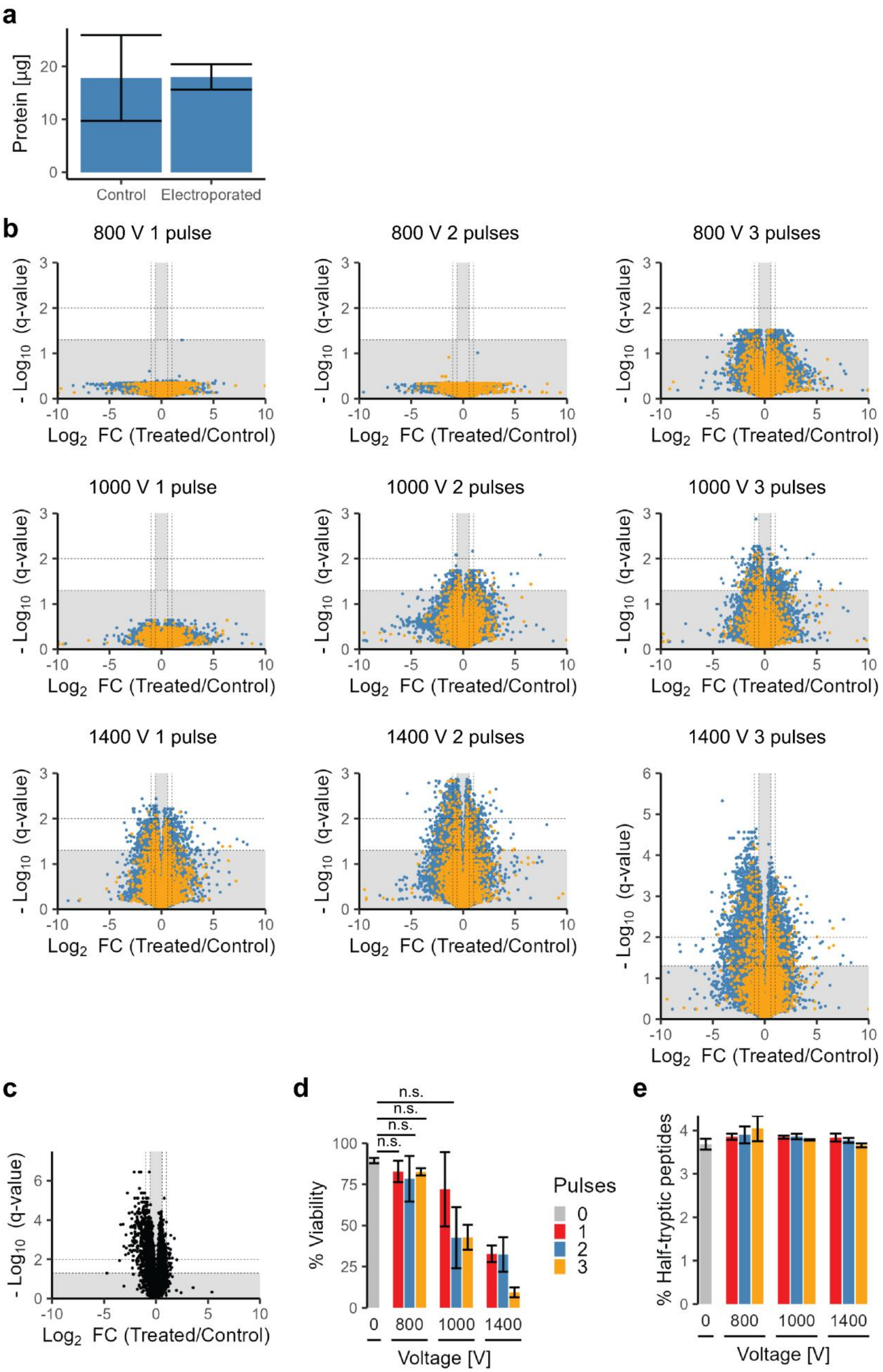
a
Protein [µg]
0
10
20
Control
Electroporated
b
800 V 1 pulse
800 V 2 pulses
800 V 3 pulses
1000 V 1 pulse
1000 V 2 pulses
1000 V 3 pulses
1400 V 1 pulse
1400 V 2 pulses
1400 V 3 pulses
- Log10 (q-value)
Log2 FC (Treated/Control)
-10
-5
0
5
10
c
- Log10 (q-value)
Log2 FC (Treated/Control)
d
n.s.
n.s.
n.s.
n.s.
% Viability
0
25
50
75
100
0
800
1000
1400
Voltage [V]
Pulses
0
1
2
3
e
% Half-tryptic peptides
0
1
2
3
4
0
800
1000
1400
Voltage [V]

***Figure S1: Effects of electroporation. (a)*** *The plot shows the amount of protein in the supernatant of 3 million HEK293 cells electroporated at 1 pulse of 1000 V, and non electroporated cells (electroporated n = 3, control n = 6).* *(**b**) The plots show peptide intensities, comparing samples with and without electroporation. Each data point represents a single peptide; half-tryptic peptides are shown in orange and fully tryptic peptides are shown in blue. The shaded gray region marks significance levels (FC ≥ 1.5, q-value < 0.05, n = 5 replicates).* *(**c**) The plot shows protein intensities at 3 pulses of 1400 V, comparing to non electroporated cells. Each data point represents a single protein. The shaded gray region marks significance levels (FC ≥ 2, q-value < 0.01, n = 5 replicates).* *(**d**) The plot shows cell viability at tested electroporation settings. Conditions that do not differ significantly from cells without electroporation are indicated with n.s. (p. adj. > 0.05).* *(**e**) The plot shows the fraction of half-tryptic peptides under the indicated conditions after in-cell LiP-MS. Colors indicate pulse number as in (d).*

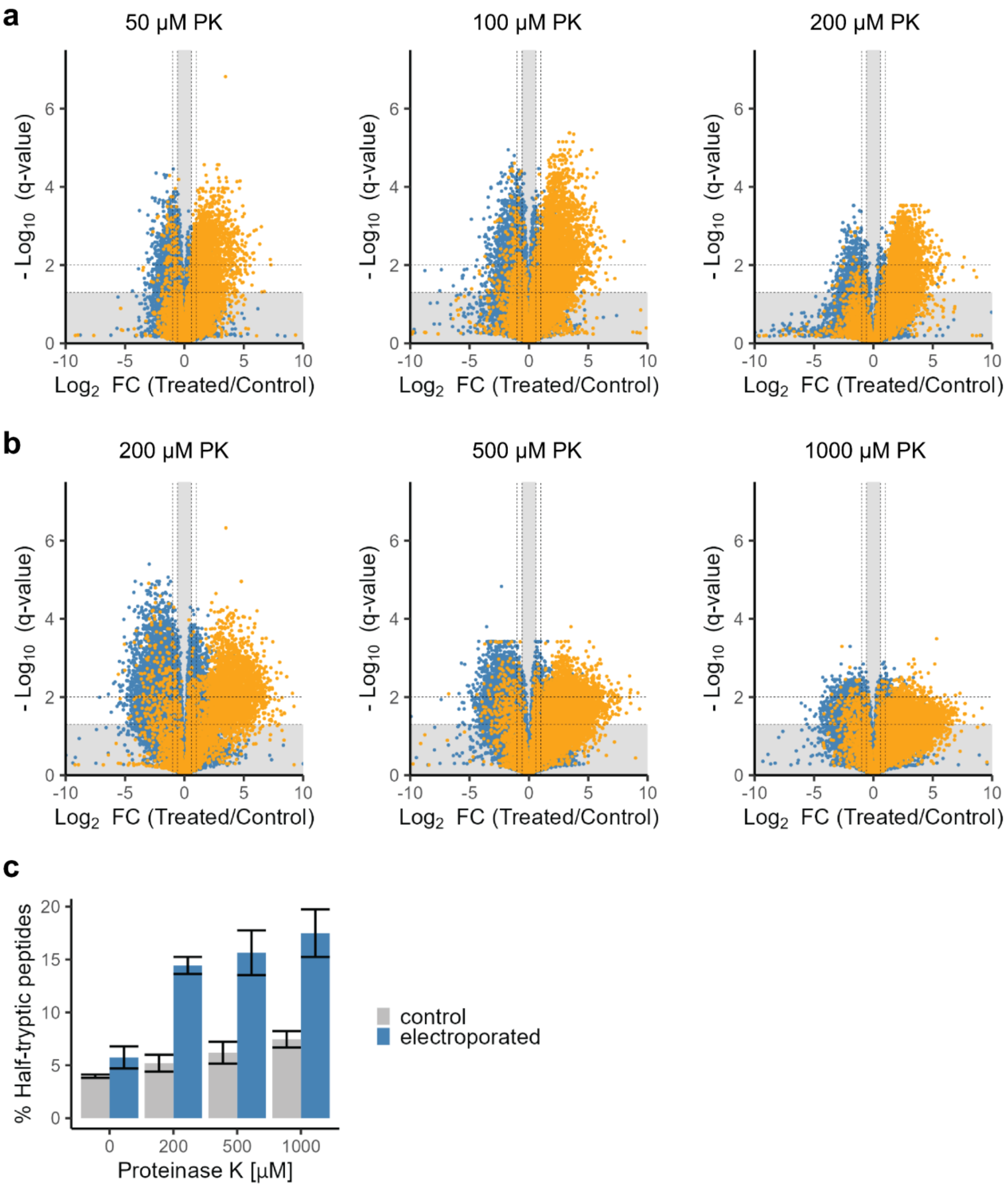


***Figure S2: Effects of PK concentration on proteolytic cleavage.*** *(**a,b**) The plots show peptide intensities after proteolytic cleavage for cells electroporated with 50 µM to 1000 µM PK, comparing samples with and without electroporation. Each data point represents a single peptide; half-tryptic peptides are shown in orange and fully tryptic peptides are shown in blue. The shaded gray region marks significance levels (FC ≥ 1.5, q-value < 0.05, n = 5 replicates). (**c**) The plot shows the fraction of half-tryptic peptides under the indicated conditions after in-cell LiP-MS.*

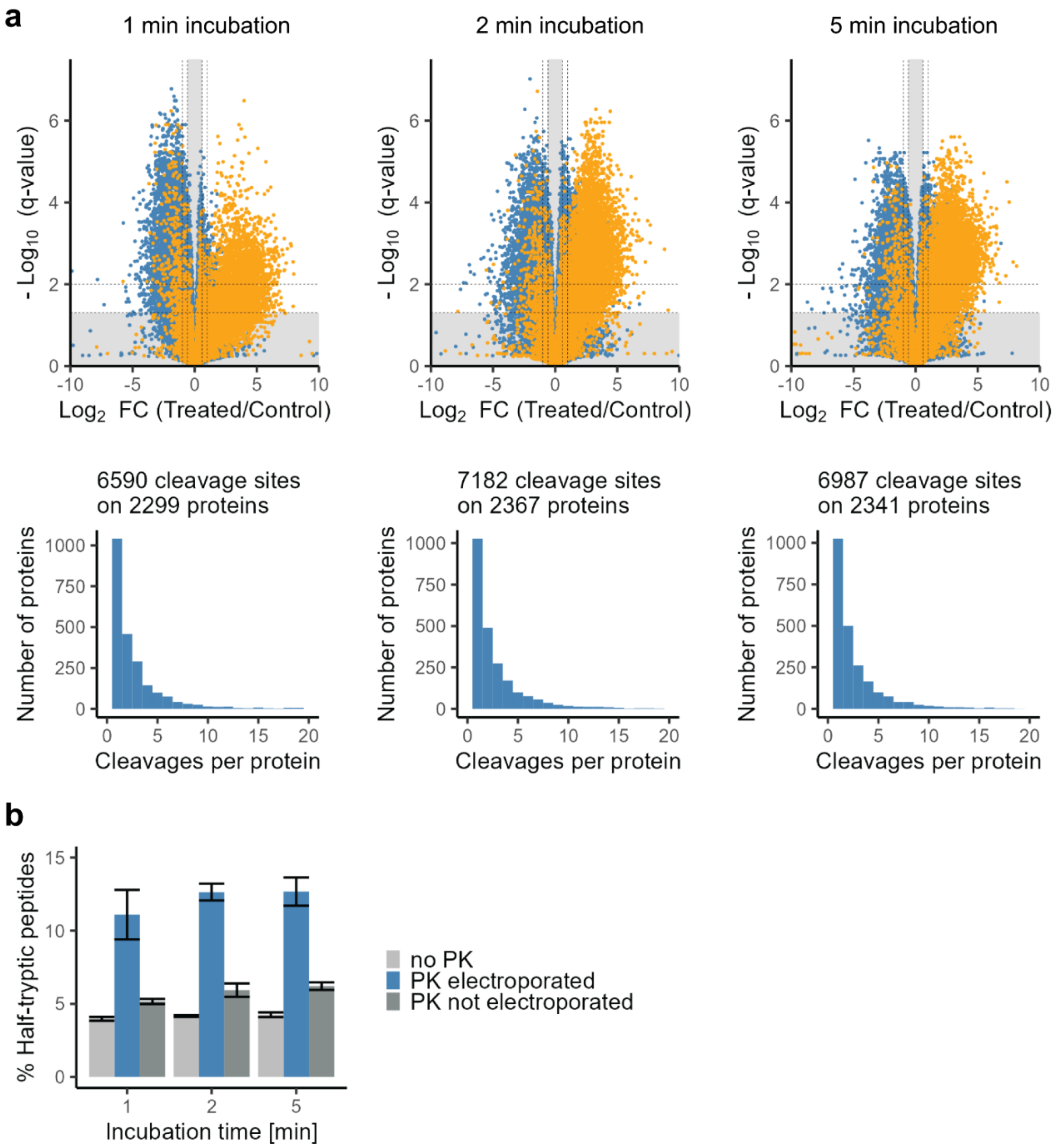


***Figure S3: Effects of incubation time upon proteolytic cleavage.*** *(**a**) The plots show peptide intensities after proteolytic cleavage for 1 min to 5 min, comparing samples with 100 µM PK with and without electroporation. Each data point represents a single peptide; half-tryptic peptides are shown in orange and fully tryptic peptides are shown in blue. The shaded gray region marks significance levels (FC ≥ 1.5, q-value < 0.05, n = 5 replicates). (**b**) The plot shows the fraction of half-tryptic peptides under the indicated conditions after in-cell LiP-MS.*

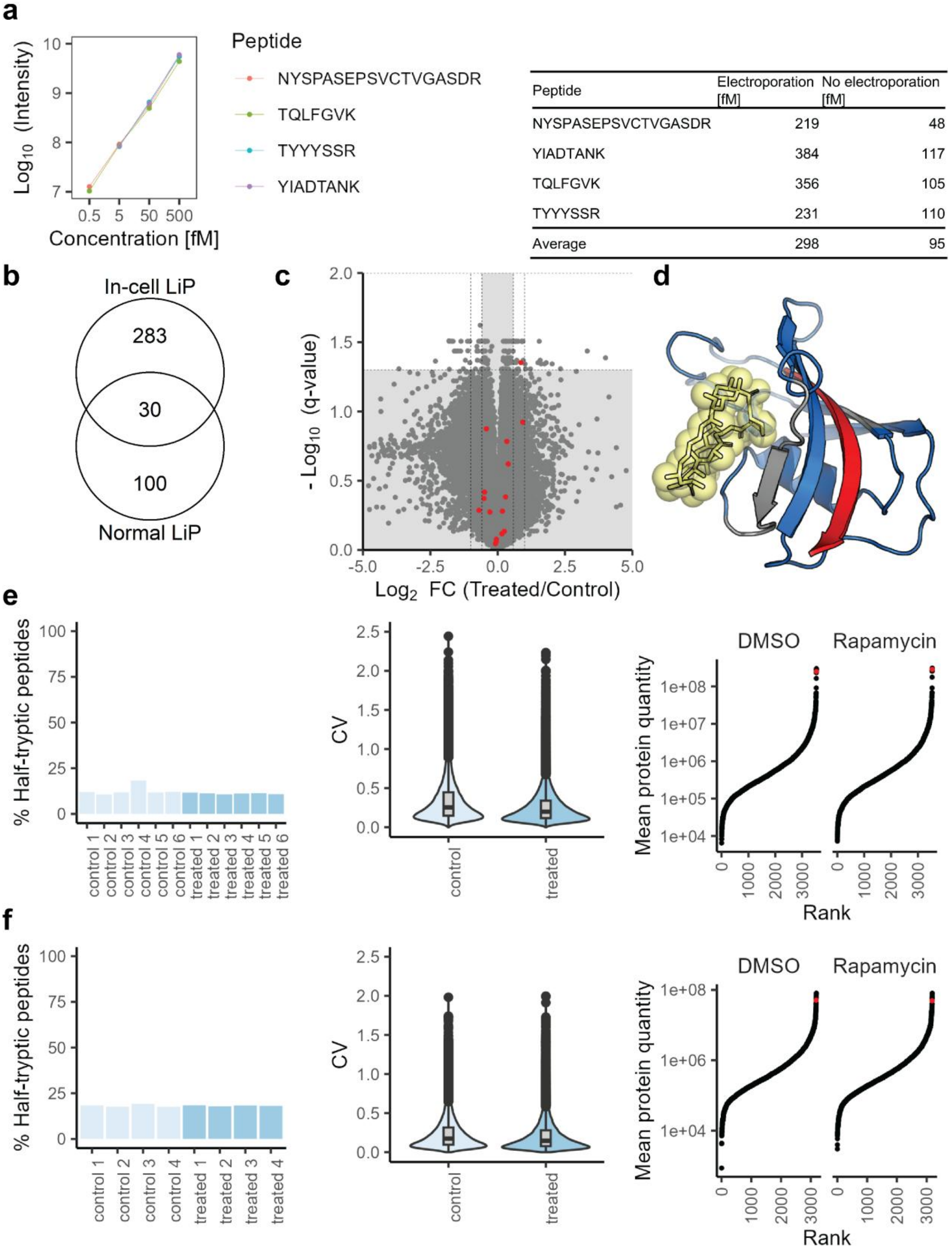


| Peptide | Electroporation [fM] | No electroporation [fM] |
|---|---|---|
| NYSPASEPSVCTVGASDR | 219 | 48 |
| YIADTANK | 384 | 117 |
| TQLFGVK | 356 | 105 |
| TYYYSSR | 231 | 110 |
| Average | 298 | 95 |

***Figure S4: Comparison of LiP-MS in cells and in native lysates. (a)** Intensity of heavy labeled peptides specific for PK spiked into HEK293 lysate at indicated concentrations. The table reports the quantity of PK peptides for in-cell LiP samples. (**b**) Overlap of proteins with significant peptide level changes for LiP of rapamycin treated lysate, and in-cell LiP of HEK293 cells treated with 100 nM rapamycin dissolved in DMSO for 5 min (FC ≥ 1.5, p-value < 0.01, n = 6 replicates for in-cell LiP, n = 4 replicates for LiP in lysate). Only proteins that are detected in both experiments are considered. (**c**) HEK283 cells treated with 10 µM rapamycin dissolved in DMSO for 10 min, compared to cells treated only with DMSO. Each data point represents a single peptide; peptides of FKBP1A are* *shown in red. The shaded gray region marks significance levels (FC ≥ 1.5, q-value < 0.05, n = 6 replicates). (**d**) Structure of rapamycin (yellow) and FKBP1A (PDB 2DG3). Regions in blue are detected by mass spectrometry, region in red is significantly changing upon treatment with rapamycin. (**e,f**) Fraction of half-tryptic peptides relative to total peptide intensity, coefficients of variation per peptide, and rank of PK intensity (red) relative to other quantified proteins. (c) Cells treated with 100 nM rapamycin for 5 min:(d) Native lysates were treated with 10 µM rapamycin for 5 min.*

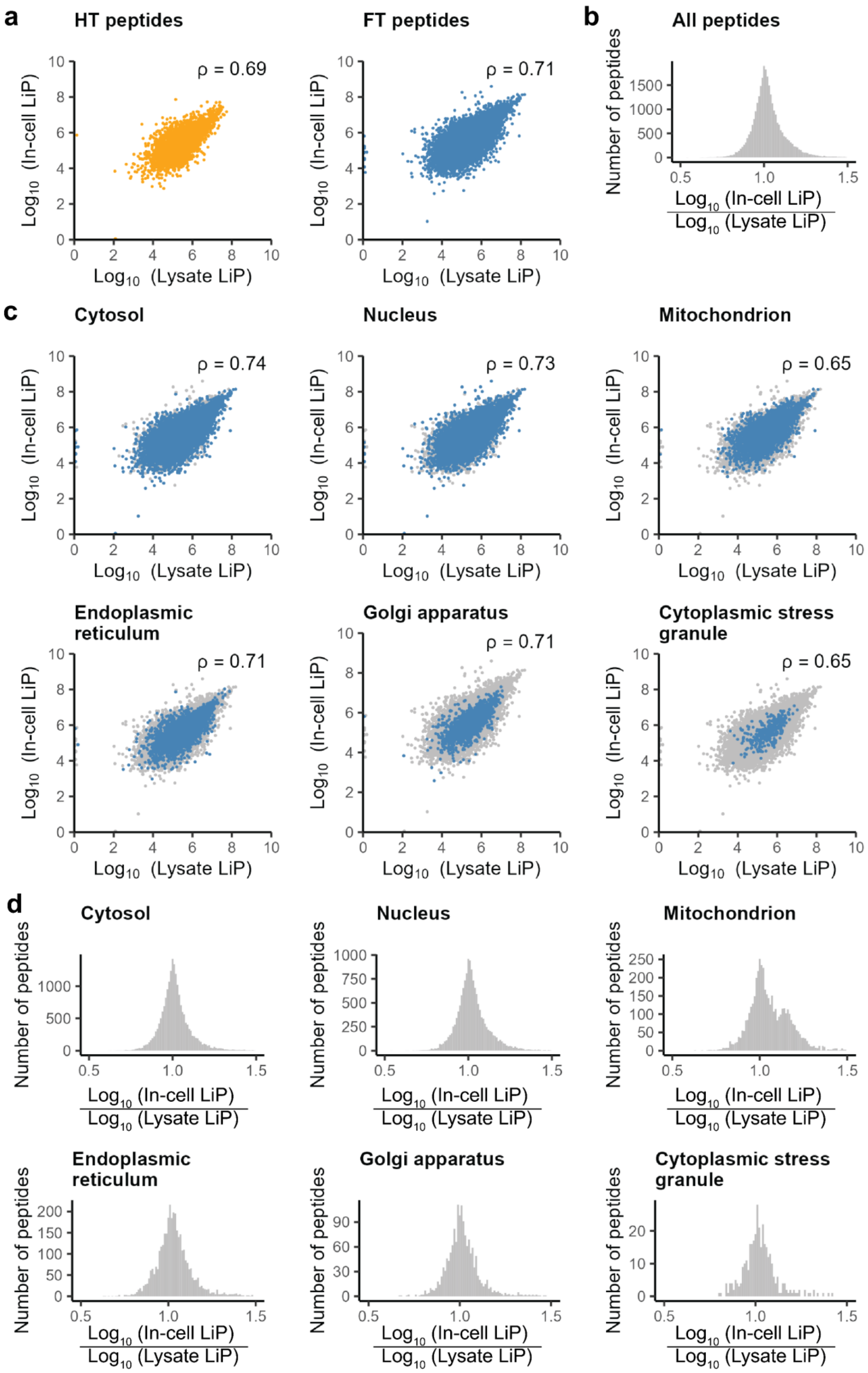
a
HT peptides
FT peptides
b
All peptides
ρ = 0.69
ρ = 0.71
Log10 (In-cell LiP)
Log10 (Lysate LiP)
Number of peptides
c
Cytosol
Nucleus
Mitochondrion
ρ = 0.74
ρ = 0.73
ρ = 0.65
Endoplasmic reticulum
Golgi apparatus
Cytoplasmic stress granule
ρ = 0.71
ρ = 0.71
ρ = 0.65
d
Cytosol
Nucleus
Mitochondrion
Endoplasmic reticulum
Golgi apparatus
Cytoplasmic stress granule

***Figure S5: Organelle coverage in LiP-MS in cells.*** *(**a**) Correlation of average peptide intensities in HEK293 cells between LiP-MS in lysate and in cells after 5 min of DMSO treatment (in-cell LiP n = 6 replicates; lysate LiP n = 4 replicates) for half-tryptic (HT) and fully tryptic (FT) peptides; ρ indicates the Pearson correlation coefficient. (**b**) Ratio of average peptide intensities in lysate and in-cell LiP. (**c**) Correlation of average peptide intensities in indicated organelles. Peptides in blue are mapping to proteins located at the indicated organelle based on gene ontology annotation in Spectronaut. Peptides from other organelles are colored grey. (d) Ratio of average peptide intensities in lysate and in-cell LiP.for indicated organelles.*

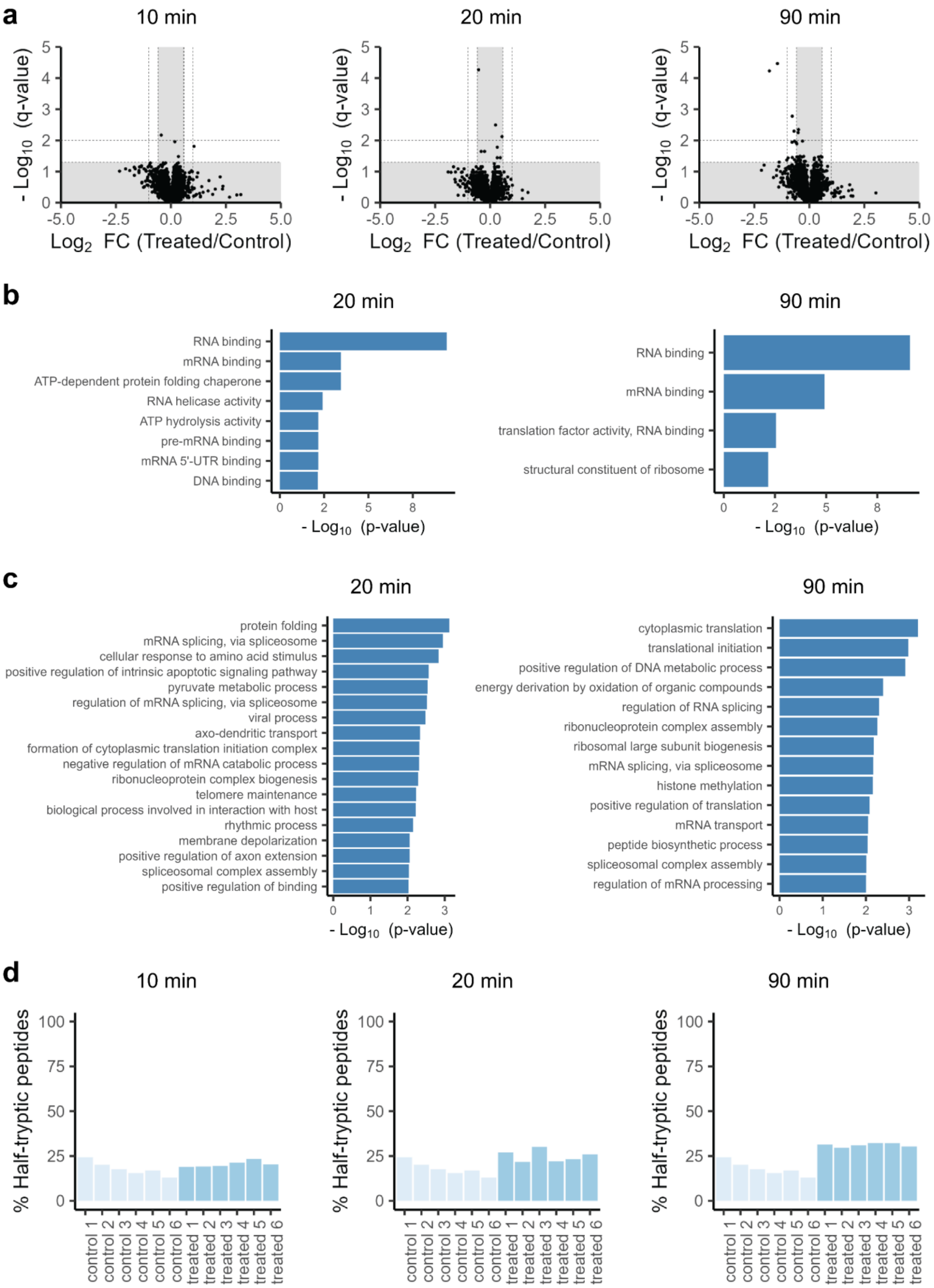


***Figure S6: Stress granule formation upon arsenite treatment.*** *(**a**) Comparison of HEK293 cells treated with sodium arsenite to untreated cells. Each data point represents a single protein. The shaded gray region marks significance levels (FC ≥ 1.5, p-value < 0.05, n = 6 replicates). (**b,c**) Gene ontology enrichment analysis of proteins with peptide-level structural changes upon arsenite treatment (p-value < 0.01). (b) Molecular function. (c) Biological*

*process (**d**) Intensity of half-tryptic peptides relative to total peptide intensity for in-cell LiP samples of arsenite treated HEK293 cells.*

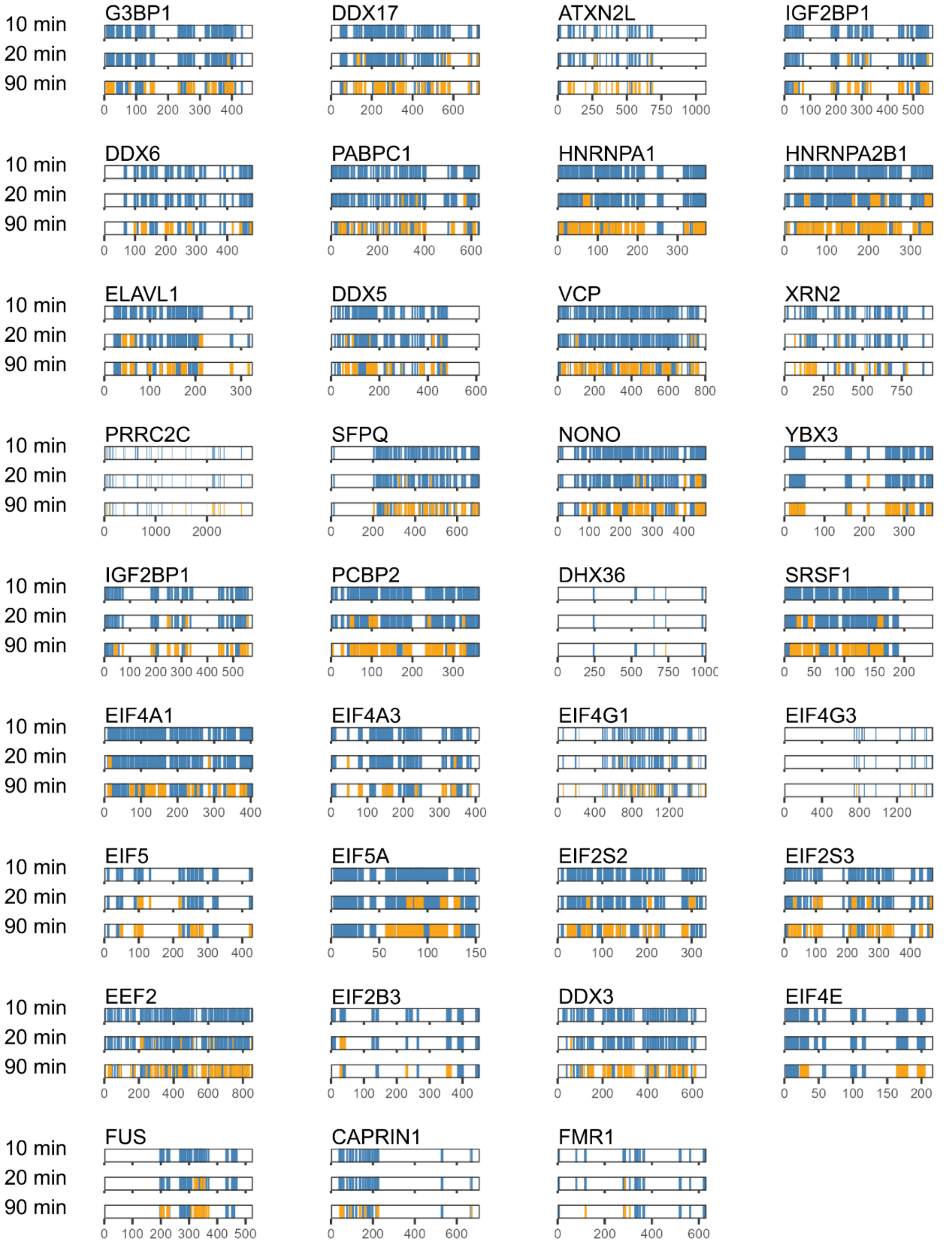


***Figure S7: Structural changes in stress granule proteins upon arsenite treatments.*** *Proteins showing structural changes upon arsenite treatment. The barcode plots depict the protein sequence, blue regions are detected by mass spectrometry. Orange regions change significantly at the indicated time (FC ≥ 1.5, p-value < 0.05, n = 6 replicates).*

### Material and Methods

#### *Cell culture conditions*

HEK293 cells (ATCC #CRL-1573) were cultured in Dulbecco's modified Eagle's medium (DMEM, Gibco Life Technologies #41965039) supplemented with 10% heat inactivated fetal bovine serum and 5% Penicillin/Streptomycin at 37 °C and 5% $CO_2$ in a humidified environment. HEK293 cells were passaged prior to confluency in phosphate-buffered saline (PBS) pH 7.4 (Gibco #10010015). Flp-In T-Rex 293 cells (Thermo Fisher R78007) were cultured under the same conditions. To measure cell viablility and concentration, cells were mixed 1:1 with trypan blue solution and analyzed in a Countess Cell Counter (Thermo Fisher Scientific).

#### *Cell transfection*

HEK239 cells were transfected at approx. 70% confluency on 150 mm culture dishes. Medium was exchanged prior to transfection. Per plate, 1200 µl jetPrime buffer were mixed with 25 µg plasmid DNA (ECFP-TevS-YPET (Gray et al., 2010), a gift from Charlie Morgan and Jason Chin; Addgene #100097), and 50 µl jetPrime reagent (Polyplus #101000046). After incubation for 10 min, the mixture was added to the cells.

#### *Fluorescence-based PK activity assay*

For cell lysate: Flp-In T-Rex 293 cells were detached from culture dishes 2-3 days after transfection, washed with PBS and lysed using a tissue grinder (DKW Life Sciences Kimble Pellet Pestle) in ten cycles of 10 s of homogenization and 1 min pause on ice. The lysate was cleared by centrifugation at 10,000 x g at 4 °C for 5 min. The supernatant was diluted to a protein concentration of 2 µg/ul, and 50 µl lysate per replicate were transferred to a 96-well plate. At the start of fluorescence measurement, 5 µl of a 100 µM PK solution (solubilized in water; in one condition electroporated with one 25 ms pulse at 1000 V), or water were added. Fluorescence of ECFP (430-10/480) and FRET to YPet (430-10/530-10) were measured using a CLARIOstar Plus plate reader (BMG Labtech).

For intact cells: HEK293 cells were detached from culture dishes 2-3 days after transfection, washed with PBS, and resuspended in PBS at a concentration of 33.3 * $10^6$ cells/ml. Prior to fluorescence analysis, 90 µl cell suspension (corresponding to $10^6$ cells) were mixed with 30 µl water or 30 µl of a 400 µM PK solution PK (solubilized in water). For electroporation, 100 µl of the solution were electroporated with one 25 ms pulse at 1000 V using a Neon Transfection System (Thermo Fisher Scientific), and transferred to a 96-well plate. Control cells without electroporation were directly transferred after mixing PK and cells. Fluorescence measurements were performed with the same settings as for cell lysates.

***Rapamycin treatment of cells***

HEK293 cells were detached from culture dishes, washed with PBS and resuspended in PBS at a concentration of 40 * $10^6$ cells/ml. For treatment, 75 µl cell suspension per replicate (corresponding to $10^6$ cells) were mixed with 15 µl rapamycin solution to a final concentration of 100 nM or 10 µM rapamycin (rapamycin dissolved in DMSO and PBS; 1% DMSO concentration on cells in the 10 µM rapamycin experiment, and 0.016% in the 100 nM rapamycin experiment). After incubation for 5 min (for 100 nM rapamycin) or 10 min (for 10 µM rapamycin), cells were processed according to the in-cell LiP-MS workflow.

***Sodium arsenite treatment of cells***

HEK293 cells were grown in separate dishes for each replicate. Sodium arsenite dissolved in water was added to the medium in a final concentration of 500 µM. After 10 min, 20 min, and 90 min, the medium was removed, cells washed in PBS, and pelleted by centrifugation at 200 x g for 1 min. Cells were resuspended in PBS. Part of the sample was snap-frozen in liquid nitrogen for analysis of protein abundances. For in-cell LiP, 90 µl containing approx. $10^6$ cells were processed according to the in-cell LiP-MS workflow.

***In-cell LiP-MS workflow***

For in-cell LiP-MS, 90 µl of treated or untreated cells (approx. $10^6$ cells) were mixed with 30 µl ice-cold PK solution (100 µM final PK concentration except in optimization experiments, PK dissolved in water). To determine the effect of PK on cells, 30 µl water were added instead. To determine the effect of electroporation independent of PK, PBS was added instead.

For electroporation, 100 µl of the cell suspension were electroporated using Neon Transfection System (Thermo Fisher Scientific). Electroporation was usually carried out with one 25 ms pulse at 1000 V, except for optimization experiments when 1-3 pulses and 800 V to 1400 V were tested. Exactly 15 seconds after electroporation, samples were heated to 37 °C, and incubated for 2 min (during optimization, 1 min to

5 min incubation were tested). For washing, 1 ml PBS pre-warmed to 37 °C was added, and cells were pelleted by centrifugation for 1 min at 4000 x g. The supernatant was removed, and cells were washed a second time with 1 ml PBS. For protease inactivation, 7 M guanidinium hydrochloride dissolved in water was added to the cells, and the samples were heated to 98 °C for at least 5 min. Samples were snap-frozen in liquid nitrogen before cell lysis.

***Cell lysis***

In-cell LiP-MS: samples were thawed on ice, and sonicated two times 30 sec using a Hielscher UP200St-G-Ultrasonic generator coupled to a UP200St with VialTweeter (optimization experiments and 10 μM rapamycin experiment: 60% cycle, 100% amplitude; stress granule experiment, 100 nM rapamycin experiment; 170 W power, 60% cycle). Protein concentration was measured using a bicinchoninic acid assay and evenly adjusted. Samples were then subjected to tryptic digestion.

Measuring protein abundances: To measure protein abundances after arsenite treatment, cell pellets were thawed on ice, and 8 M urea buffer supplemented with protease inhibitor cocktail (Roche) was added. Cells were sonicated like the in-cell LiP-MS samples of the stress granule experiment, protein concentration was measured using a bicinchoninic acid assay and evenly adjusted. Samples were then subjected to tryptic digestion.

Classical LiP-MS: HEK293 cells were detached from culture dishes, washed with PBS and resuspended in a buffer consisting of 1 mM MgCl2, 150 mM KCl, 100 mM HEPES pH 7.4. Cells were lysed using a tissue grinder (DKW Life Sciences Kimble Pellet Pestle) in ten cycles of 10 s of homogenization and 1 min pause on ice. The lysate was cleared by centrifugation at 10000 x g at 4 °C for 15 min. The supernatant was diluted to a protein concentration of 2 μg / μl and treated with rapamycin.

***Classical LiP-MS workflow of rapamycin treatment***

For classical LiP-MS, 50 μl native lysate of 2 μg / μl protein were incubated with 1 μl rapamycin (dissolved in DMSO), or 1 μl DMSO for 5 min at 25 °C. To start limited proteolysis, 5 μl PK (0.2 μg / μl in water) were added for 5 min. For protease inactivation, samples were heated to 99 °C for 5 min, then cooled to 4 °C for 5 min. Finally, an equal volume of a 10% sodium deoxycholate solution was added. Samples were subjected to tryptic digestion.

***Tryptic digestion***

Disulfide bonds were reduced by incubation with 5 mM tris(2-carboxyethyl)phosphine hydrochloride) (TCEP) shaking for 40 min at 37 °C. Subsequently, samples incubated 30 min in the dark at room temperature with 40 mM. Samples were then diluted using 0.1 M ammonium bicarbonate (Sigma Aldrich) to achieve a final guanidinium chloride concentration of 0.5 M, or 1% sodium deoxycholate, or <2 M urea. Proteins were digested with Lys-C (FUJIFILM Wako Pure Chemical Corporation) and trypsin (Promega) at an enzyme substrate ratio of 1:100 shaking overnight at 37 °C. The tryptic digest was stopped by adding formic acid to a final pH of approx. 2.

***C18-Cleanup and MS sample preparation***

A 96-well C18 Micro-Spin plate (The Nest Group, 40-400 µg) was washed with 200 µl methanol, followed by 100 µl buffer B (50% acetonitrile (ACN), 0.1% FA), and 2 x 200 µl buffer A (0.1% FA). Samples were loaded on the plate and washed with 3 x 200 µl buffer A by centrifugation at 1000 x g for 1 min. Peptides were eluted with 3 x 100 µl buffer B and dried in a vacuum concentrator. Samples were resuspended in buffer A supplemented with iRT peptides (Biognosys) and analyzed by mass spectrometry.

***Absolute quantification of PK***

Four synthetic peptides (AQUA Ultimate Heavy, lysines and arginines stable isotope-labeled, Thermo Scientific) that are proteotypic for PK were selected for absolute quantification (TQLFGV(K), TYYYSS(R), YIADTAN(K),NYSPASEPSV[C-carbamidomethyl]TVGASD(R)). Heavy labeled peptides were added at a final concentration of 500 fmol/µl before mass spectrometry analysis to an in-cell LiP sample electroporated previously with 100 µM PK. Peptides were also added to a control sample to which PK was added but not electroporated. For calibration, heavy peptides were spiked into a lysate containing no PK (peptide concentrations 500 fmol / µl, 50 fmol / µl, 5 fmol / µl, 500 amol / µl). Samples were measured by PRM. Peptide intensity was determined by summing up the area of the five most intense fragment ions. A linear model was fitted separately for each peptide to $\log_{10}$-transformed peptide intensities and concentrations of the calibration samples. The linear model was used to quantify the corresponding light PK peptides in the samples of interest. Finally, PK quantity was calculated as the mean of the four determined peptide quantities.

***Liquid Chromatography and mass spectrometry data acquisition***

DIA: The in-cell LiP-MS samples of 100 nM rapamycin and arsenite were injected into a Orbitrap Fusion Lumos Tribrid mass spectrometer equipped with a Waters nanoAQUITY UPLC System using a linear gradient from 3-35% B (Eluent A: 0.1% formic acid, Eluent B: 99.9% acetonitrile, 0.1% formic acid) over 120 minutes at a flowrate of 300 nL / min. The MS1 scans ranged from 350 to 1400 m/z with an Orbitrap resolution of 120000 and an AGC target of 50% or 100 ms injection time. The DIA method consisted of 41 variable width windows with 1 m/z overlap. Fragments were generated by high energy collision induced dissociation (HCD) using a fixed collision energy of 28%. MS2-DIA spectra were acquired over a scan range of 200 to 1800 m/z with an Orbitrap resolution of 30000 and 200% AGC target or 54 ms injection time.

DIA: The in-cell LiP-MS samples of 10 µM rapamycin were injected into a Q Exactive Plus Hybrid Quadrupole-Orbitrap mass spectrometer equipped with an EASY-nLC 1000 system (Thermo Fisher Scientific). Separation of peptides was conducted on a 40 cm x 0.75 mm i.d. column (New Objective, PF360-75-10-N-5) which was in house packed with 3 µm beads (Dr. Maisch Reprosil-Pur 120). Separation followed a linear gradient of 3% - 30% buffer B over a total run time of 120 minutes, followed by 5 min with an isocratic constant concentration of 90% buffer B (Eluent A: 0.1% formic acid, Eluent B: 95% acetonitrile, 0.1% formic acid). The flow rate was kept at 300 nl / min. The MS1 scans ranged from 350 to 1500 m/z with an Orbitrap resolution of 70,000 and an AGC target of 3e6 or 120 ms injection time. The DIA method consisted of 20 variable width windows with 1 m/z overlap. Fragments were generated by high energy collision induced dissociation (HCD. MS2-DIA spectra were acquired over a scan range of 200 to 1800 m/z with an Orbitrap resolution of 35,000 and 1e6 AGC target.

DIA: Samples for optimization of electroporation settings were injected into a Orbitrap Eclipse Tribrid mass spectrometer equipped with a Waters nanoAQUITY UPLC System using a linear gradient from 3-30% B (Eluent A: 0.1% formic acid, Eluent B: 95% acetonitrile, 0.1% formic acid) over 120 minutes at a flowrate of 300 nL / min. The MS1 scans ranged from 350 to 1400 m/z with an Orbitrap resolution of 120,000 and an AGC target of 800,000. The DIA method consisted of 41 variable width windows with 1 m/z overlap. Fragments were generated by high energy collision induced dissociation (HCD. MS2-DIA spectra were acquired over a scan range of 150 to 2000 m/z with an Orbitrap resolution of 30,000 and 50,000 AGC target.

PRM: Samples were injected into a Orbitrap Fusion Lumos Tribrid mass spectrometer equipped with a Waters nanoAQUITY UPLC System using a linear gradient from 3-35% B (Eluent A: 0.1% formic acid, Eluent B: 99.9% acetonitrile, 0.1% formic acid) over 120 minutes at a flowrate of 300 nL / min. The isolation window was 1.2 m/z. Fragments were generated by high energy collision induced dissociation (HCD) using a fixed collision energy of 28%. MS2 spectra were acquired over a scan range of 200 to 1800 m/z with an Orbitrap resolution of 60000 and 1000% AGC target or 118 ms injection time.

***Search engine***

The data was searched using Spectronaut 17 (Biognosys), with libraries built from the DIA files. The digestion type was set to semi-specific and peptide minimum length was adjusted to 6. The data was searched against the human reviewed UniProt database and the sequence of PK (https://www.uniprot.org, retrieved on April 16, 2023). Data analysis was performed with Spectronaut via a classical DIA analysis using a targeted library-based search with the following modifications: a single-hit was defined by modified sequence, and single-hit proteins were excluded. Quant 2.0 was selected for label-free protein quantification, samples were normalized globally on the median, and PTM localization was selected.

***Statistical analysis***

Only peptides with a quantity <1 were considered for statistical analysis using R version 4.2.2. A Welch’s t-test was used for statistical analysis. P-values were adjusted to multiple testing using the Benjamini Hochberg method.

***Gene ontology enrichment analysis***

For gene ontology enrichment analysis, the topGP package (version 2.50.0) was used in R (https://git.bioconductor.org/packages/topGO). Gene ontology annotations were obtained from Uniprot (https://www.uniprot.org, retrieved on May 17, 2022). Enrichment analysis was performed with the Fishers exact test using the elim algorithm in topGO. Significance was set at the p-value level < 0.01.

***Peptide mapping on AlphaFold2 structures***

Peptides were mapped onto structures predicted by Alphafold v4 using PyMOL (version 2.5.4).

***Immunostaining and circularity measurements***

HEK293 cells were seeded on 8 well µ-Slides with ibiTreat surface (ibidi #80806) at 25000 cells per well. After 24 h 500 µM sodium arsenite (Pfaltz & Bauer #7784-46-5) was added to the cells. After 90 mins cells were fixed for 10 min with 4% PFA (Electron Microscopy sciences #15714) in PBS. After washing twice with PBS the cells were permeabilized for 10 min in PBS with 0.5% Triton X-100 (Sigma-Aldrich #T8787). Cells were washed twice with PBS and then incubated simultaneously with primary antibody, secondary antibody and 1:1000 Hoechst 33258 (Molecular Probes #H-3569) for 1 h in PBS with 0.1% Tween 20 (Sigma-Aldrich #P1379). Cells were washed afterwards 2 times in PBS with 0.1% Tween 20 and kept in PBS for imaging.

To determine the circularity of nuclear speckles, a representative field of view of each replicate was taken and thresholded in imageJ to create a mask of the nuclear speckles that matches the observed speckles by eye. Circularity was then measured via the analyse particles function.

Antibodies used:

| Name | Supplier | Catalog# | Concentration |
|---|---|---|---|
| Anti-G3BP1 | Santa Cruz Biotechnology | sc-365338 | 1:1000 |
| Anti-SC35 | Sigma-Aldrich | S4045 | 1:2000 |
| Alexa Fluor 488, anti-Mouse | Life technologies | A-11029 | 1:1000 |

### References

Gray, D. C., Mahrus, S., & Wells, J. A. (2010). Activation of Specific Apoptotic Caspases with an Engineered Small-Molecule-Activated Protease. *Cell*, *142*(4), 637–646. https://doi.org/10.1016/j.cell.2010.07.014
